# The genetic origins and impacts of historical Papuan migrations into Wallacea

**DOI:** 10.1101/2024.06.02.597070

**Authors:** Gludhug A. Purnomo, Shimona Kealy, Sue O’Connor, Antoinette Schapper, Ben Shaw, Bastien Llamas, Joao C. Teixeira, Herawati Sudoyo, Raymond Tobler

## Abstract

The tropical archipelago of Wallacea was first settled by anatomically modern humans (AMH) by 50 thousand years ago (kya), with descendent populations thought to have remained genetically isolated prior to the arrival of Austronesian seafarers around 3.5 kya. Modern Wallaceans exhibit a longitudinal countergradient of Papuan- and Asian-related ancestries widely considered as evidence for mixing between local populations and Austronesian seafarers, though converging multidisciplinary evidence suggests that the Papuan-related component instead comes primarily from back-migrations from New Guinea. Here, we reconstruct Wallacean population genetic history using more than 250 newly reported genomes from 12 Wallacean and three West Papuan populations and confirm that the vast majority of Papuan-related ancestry in Wallacea (∼75–100%) comes from prehistoric migrations originating in New Guinea and only a minor fraction is attributable to the founding AMH settlers. Mixing between Papuan and local Wallacean lineages appear to have been confined to the western and central parts of the archipelago and likely occurred contemporaneously with the widespread introduction of genes from Austronesian seafarers—which now comprise between ∼40–85% of modern Wallacean ancestry—though dating historical admixture events remains challenging due to mixing continuing into the Historical Period. In conjunction with archaeological and linguistic records, our findings point to a dynamic Wallacean population history that was profoundly reshaped by the spread of Papuan genes, languages, and culture in the past 3,500 years.

## INTRODUCTION

The Wallacean archipelago is a world-renowned biodiversity hotspot lying between the former Pleistocene continents of Sunda (connecting the Malay Peninsula, Sumatra, Borneo, Java, and Bali) to the west and Sahul (comprising Australia, Tasmania, New Guinea, Misool and the Aru Islands) to the south and east. Since their orogenic and volcanic origins in the Cenozoic, each of the >1,000 Wallacean islands have remained separated from their continental neighbours, and one another, by a series of oceanic straits (1). These passages have presented persistent dispersal barriers for historical migrants, resulting in the emergence of a transitional biographical zone bounded by the Wallace and Lydekker Lines to the west and east, respectively (2) (**Fig. 1**). Despite these obstacles, lithic and fossil records attest to multiple hominin groups successfully crossing Wallace’s Line in the late Pleistocene (3–5), with AMH arriving by at least 50 kya as part of a wider radiation of our species beyond Africa that culminated in the peopling of Sahul around the same time (6–8).

**Figure 1.**
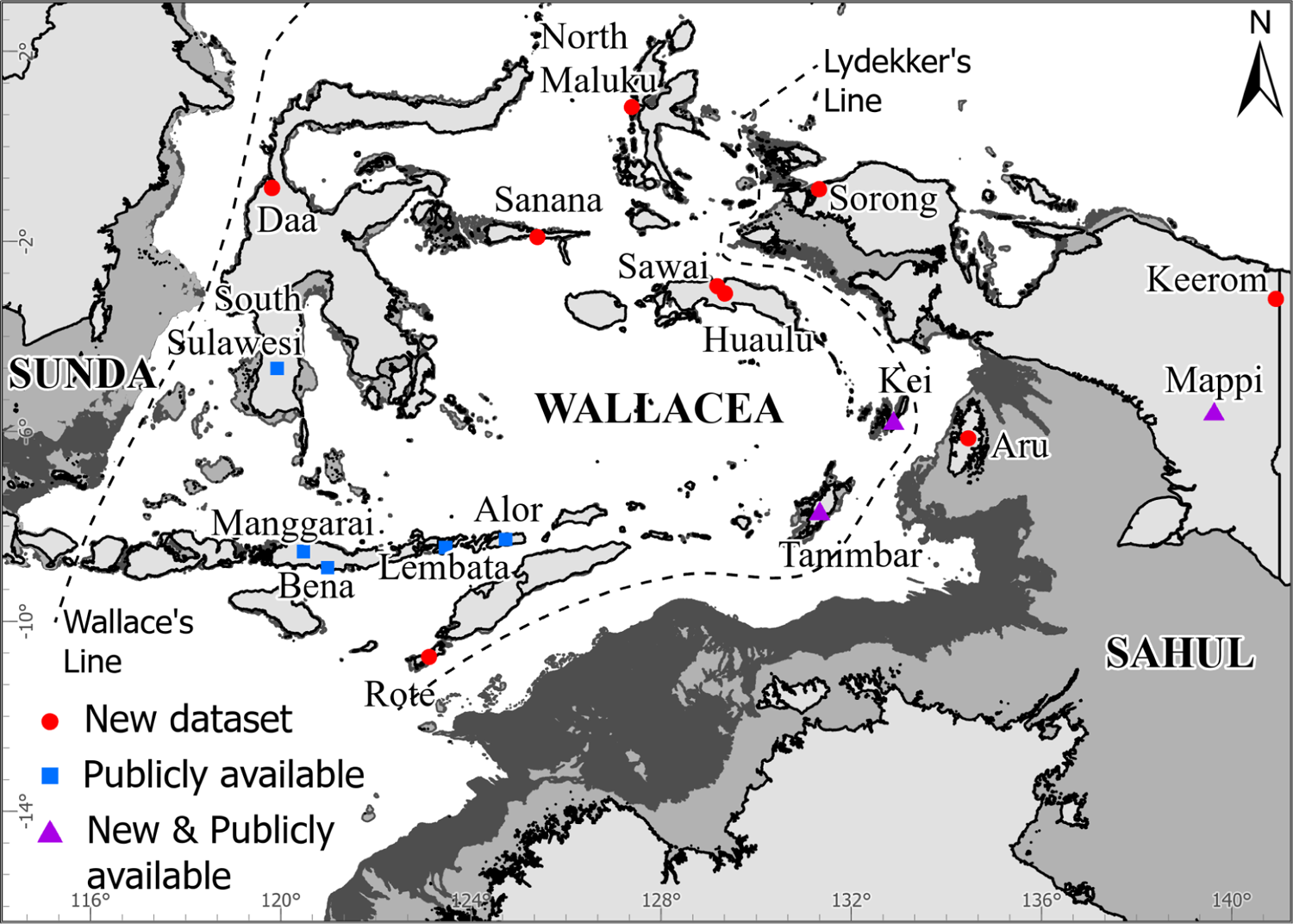
Map of the Wallacean archipelago and sampling locations. Contemporary coastlines are demarcated by black lines, with historical coastlines at ∼60–50 kya (−65m below present sea level, bpsl) and LGM (−130m bpsl) indicated by successively darker grey shading (bathymetric data sourced from GEBCO 2021 dataset), when lower sea levels exposed the neighbouring continents of Sunda (comprising the Malay Peninsula, Sumatra, Borneo, Java, and Bali) and Sahul (comprising Australia, New Guinea, Misool and the Aru Islands). Coloured points indicate the sampling locations of newly reported and previously published Wallacean and West Papuan genomes used in this study (see key). Newly reported samples come from Kei (*n* = 20), Aru (*n* = 23), Tanimbar (*n* = 22), Huaulu (*n* = 20), Sawai (n = 7), North Maluku (*n* = 30), Sanana (*n* = 19), Daa (*n* = 22), Rote-Ndao (n = 29), Keerom (*n* = 26), Mappi (*n* = 11), and Sorong (*n* = 27). A complete list of all samples and associated metadata is provided in Table S1.

The significance of Wallacea as a hominin evolutionary hub (9, 10) and gateway for historical human migrations to Sahul has motivated a growing number of studies exploring the human occupation of the region. These studies have revealed a dynamic past involving movement, mixing, and trade among multiple human groups (11), resulting in parallel counter-gradients of Papuan- and Asian-related genetic ancestries (12–15) and linguistic forms (16) across modern Wallacea. However, while multiple lines of evidence attest to Austronesian seafarers being the primary source of Asian-related genes and languages in Wallacea from ∼3.5 kya onwards (17), doubt persists about whether Papuan-related ancestry is evidence for descent from the founding AMH settlers or the result of subsequent arrivals from New Guinea (18).

At present, perhaps the best evidence for genetic continuity following the initial settlement of Wallacea comes from a middle-Holocene Toalean forager who lived in Southern Sulawesi ∼7.2 kya, with approximately half of their ancestry coming from a lineage that diverges from Near Oceanians (i.e., modern Aboriginal Australians and Melanesians) around the same time that these two groups also separate (19). However, Toalean forager ancestry was not detected in individuals living in Sulawesi and a handful of Eastern Wallacean islands from ∼2.5 kya to the present (20), with all examined individuals instead being more closely related to modern Papuan and Asian groups. Evidence for the retention of founder AMH ancestry is also scarce in mitogenome and Y-chromosome datasets, with two recent phylogenetic studies finding that Wallacean lineages only rarely appear in basal phylogenetic positions relative to Australo-Papuans and are far more likely to occur nested within Papuan clades (21, 22). Accordingly, mounting genetic evidence suggests that a significant fraction of Papuan-related ancestry in Wallacea may derive from back-migrations from New Guinea rather than wholly attesting to descent from AMH founders.

Growing archaeological and linguistic evidence also supports a central role for Papuan people in shaping Wallacean history. Technologies (23), art (24, 25), fauna (26), and the recently proposed Greater West Bomberai (GWB) language family spoken in eastern Wallacea are all thought to be introductions from West Papua (27). The timings of putative Papuan back-migrations inferred from mitogenomes also bear close chronological concordance to the emergence of novel maritime technologies and inter-island trading networks in eastern Wallacea at the end of the LGM (∼15 kya; (24)), with a second pulse occurring around the time that Austronesian seafarers started spreading across the archipelago. However, most details of the underlying historical Papuan migration—including the identity of the source populations and their connection to the arrival of Austronesian migrants and to broader demographic changes occurring in New Guinea (28)—remain unknown. These fundamental knowledge gaps highlight the need for broader population genomic surveys across modern Wallacea to illuminate its broad geo-genetic structure and investigate the origins and impacts of these enigmatic Papuan migrations.

## RESULTS

### Comprehensive Wallacean and West Papuan population genomic dataset

To address these outstanding questions in Wallacean genetic history, we applied shallow shotgun sequencing (average coverage ∼2.3x, range 0.9x to 4.6x) to 256 individual genomes collected from 12 different communities across Wallacea and three from West Papua—comprising around 30 different ethnic and dialect groups (**Fig. 1, Table S1**)—following informed consent protocols established by the Indonesian Genome Diversity Project (12, 29). These genomes were supplemented by 161 previously published high coverage Indonesian and Melanesian genomes (12, 29), increasing the total number of Wallacean populations to 14 and providing good representation along the two major migratory routes—i.e., the Northern Route (with samples from Sulawesi [South Sulawesi, *N* = 12, Daa, *N* = 21], Sanana [*N* = 19], North Moluccas [*N* = 30], Seram [Sawai, *N* = 7, Huaulu, N = 20], Kei [*N* = 24], Tanimbar [*N* = 28]) and the Southern Route (samples from Flores [Bena, *N* = 11, Mangarrai, *N* = 34], Rote [*N* = 28], Lembata [*N* = 7], and Alor [*N* = 5])—proposed to have been traversed by the first AMH settlers of Sahul (sample sizes and ethnographic subgroupings in square brackets; see **Fig. 1 and Table S1**).

Our sampling also includes the first reported data from the eastern Wallacean island of Aru [*N* = 27], which was formerly situated on the Sahul coastline before becoming separated by rising sea levels around 10 kya (30), and also some of the first reported West Papuan population genetic datasets—including individuals from Keerom [*N* = 26] and Mappi [*N* = 18] in the central-north and southwestern New Guinea, respectively, and Sorong [*N* = 27] in the western Bird’s Head region. These groups facilitate investigation of the poorly defined West Papua region and Papuan genetic history more generally, and capture potential entry points into Sahul in addition to possible historical exit points from New Guinea into Wallacea.

For all newly reported Wallacean and West Papuan genomes, genotypes were called for each individual using GLIMPSE (v1.1; (31)), which draws upon genetic information from a phased genetic reference panel (i.e., the Human Genome Diversity Panel, HGDP; (32)) and from the target samples to provide accurate imputed genotype calls (31). To assess the quality of our genotype calls, we resequenced eight samples to high coverage (average ∼28x) and generated high accuracy genotype calls for each, which were subsequently compared to the complementary low coverage calls (**see SI Methods**). These analyses indicate that the imputed genotype data provide accurate and robust results when applying the population genomic inference methods employed in the current study (**Fig. S1;** full details provided in ref. (33)).

### Papuan- and Asian-related ancestries dominate the modern Wallacean genetic landscape

To place the Wallacean populations within a broader global context and uncover genetic structure within Wallacea and West Papua, we combined our Indonesian and Melanesian samples with 438 additional high coverage genomes from global human populations (13, 34, 35). After removing 11 closely related individuals (1st and 2nd-degree relatives; **see Methods**), we submitted the resulting dataset of 844 individuals to Principal Components Analysis (PCA; performed using smartsnp v1.1.0; (36)) and genetic ancestry decomposition (performed with ADMIXTURE v1.30; (37)) (**see Methods**).

The distinctive dual ancestry countergradient reported in previous genetic studies of Wallacea (12–15) is also a defining feature in both the PCA and ancestry decomposition analysis of our considerably larger grouping of 270 Wallacean individuals (**Figs. 2A and S2**). Wallacean ancestry is effectively decomposed into two genetic sources in all ADMIXTURE models from *K* = 3 to *K* = 8 components, which find their highest representation in modern genetic proxies for the original Austronesian-speaking seafarers (i.e., an Indigenous Taiwanese group comprising Ami and Atayal; denoted by rich blue components in **Figs. 2A and S2**) or in mainland Papuan populations (i.e. teal components in **Figs. 2A and S2**). Asian-related ancestry shows a general increase from east to west that is countered by decreasing Papuan-related ancestry, which is highest in eastern Wallacea (i.e., ∼60% in Aru) and becomes negligible in populations living in western Indonesian regions that were formerly part of Sunda (e.g., Java, Borneo, Sumatra, Mentawai, Nias; WSE group in **Fig. 2**). The familiar counter-clinal distribution of Asian- and Papuan-related ancesties is also evident along the second dimension of the PCA (**Figs. 2B and S3**), with populations from eastern Wallacea being situated closer to Papuan groups at one end of the cline, and those from the western part of the archipelago lying nearest to Southeast Asian and Indigenous Taiwanese groups at the opposite end. Notably, this pattern is closely reproduced along the first dimension of the PCA formed by projecting Wallacean and west Indonesian individuals onto a PCA space limited to genetic diversity from populations from Southeast Asia, Taiwan, and PNG (**Fig. 2C and S4**), confirming that Wallacean ancestry can be effectively modelled as a countergradient of Asian- and Papuan-related ancestries.

**Figure 2.**
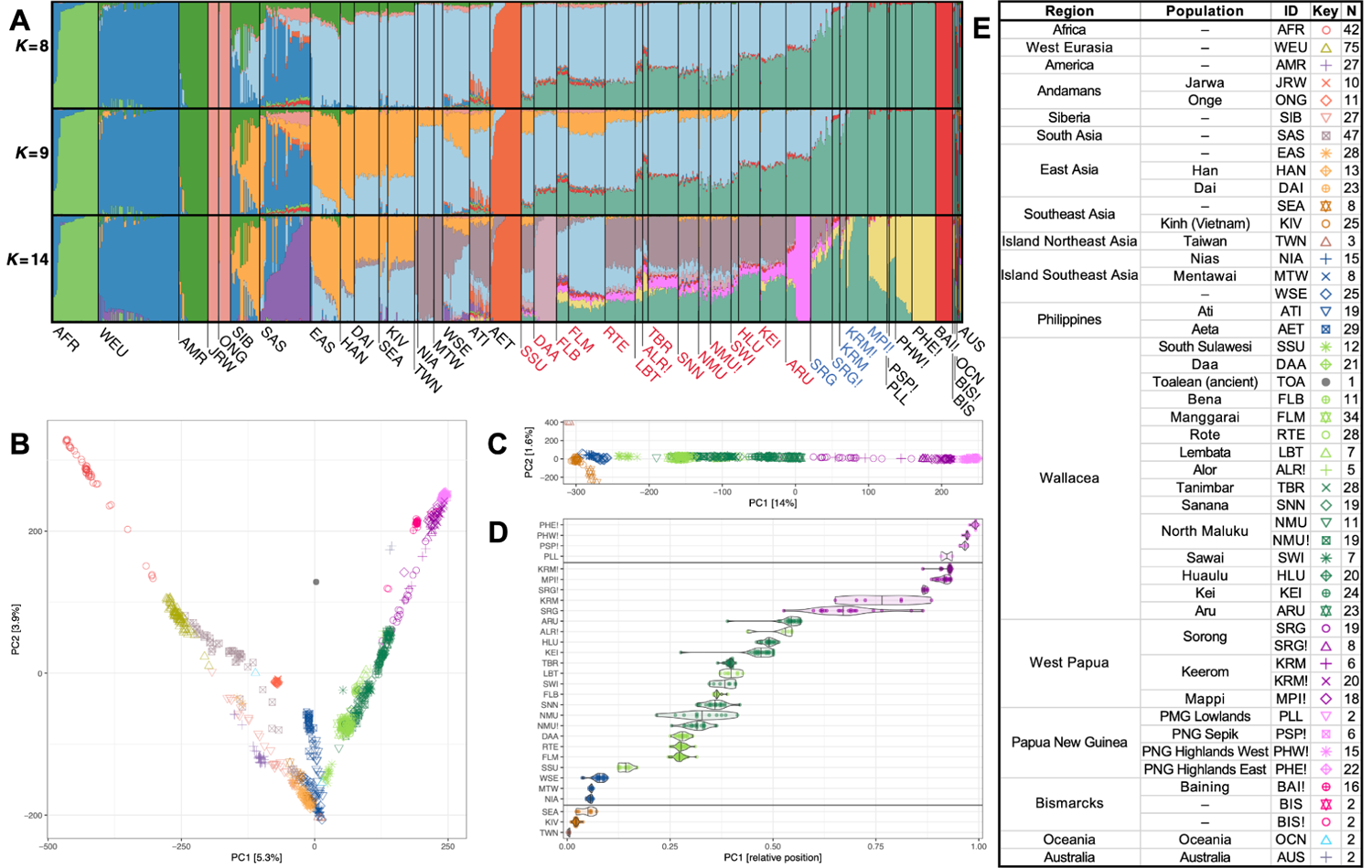
Countergradients of Asian- and Papuan-related ancestry in Wallacea. **A.** Fractional contributions of ancestry (different coloured bars) from genetically distinct source groups (number of groups, *K,* shown on left side of plot) in a global dataset of 855 individuals. Populations in the Wallacean archipelago (red text) show increasing amounts of Asian-related ancestry, and declining Papuan-related ancestry, running from eastern Indonesia to West Papua (blue text). Note that the island of Aru is physically located on Sahul shelf but is included as a Wallacean grouping throughout for convenience, and this post-hoc classification does not impact any of our analyses. Wallacean and Melanesian groups are further separated into linguistic subgroupings (exclamation marks indicate a Papuan language is spoken, remaining groups are Austronesian language speakers). Counterclinal patterns are also evident in the 2nd dimension of a PCA performed on the same global dataset (**B**) and following projection of Wallacean and West Papuan individuals onto a PCA (**C**) generated using select Asian and Papuan individuals (population symbols for both PCAs shown in key in panel E). **D.** Violin plot showing distribution of individuals in the first dimension of the PCA in panel C, with populations ordered by median position on y axis. **E.** Geographical summary of global population genomic datasets used in this study, including abbreviated names (ID), symbols used in panels B and C, and sample sizes (N) (additional details provided in **Table S1**).

Extending the number of ancestry components from nine to 15 sources in our ADMIXTURE models revealed additional genetic structuring across Wallacean populations, though many components capture structure among large Wallacean groups that are not readily informative of historical regional migrations or mixing events (**Figs. 2A and S2**). A key exception comes from the best-fitting ADMIXTURE model (*K* = 9 ancestry components), where a source that is most prominent in East and Southeast Asia (i.e., denoted by the mango components in **Figs. 2A and S2**) appears at low to moderate levels in most Wallacean groups (population means ∼1% to 30%). This ancestry source tends to be less common in eastern Wallacea and almost entirely absent among Papuan groups, echoing findings from previous Indonesian population genomic studies that infer the presence of an Southeast Asian-related ancestry source in parts of Wallacea that is genetically distinct from proxies for Austronesian seafarers (12, 20). Recent paleogenomic evidence suggests that these ancient migrants are related to groups from Thailand and Laos and were actively spreading through Wallacea between 2.5 to 1 kya (20).

Internal structuring among Papuan groups becomes apparent at *K* = 14 ancestry components (mustard component in **Figs. 2A and S2**), with this new component being most strongly associated with populations from the Eastern Highland region of PNG and the other Papuan ancestry source now being most abundant among the West Papuan Keerom population (**Figs. 2A and S2**). Notably, this Eastern Highland associated ancestry is rare or absent among groups in West Papua and most of Wallacea, being most notable in populations from the Lesser Sunda islands (i.e., Flores, Lembata, Alor). These patterns suggest that multiple sources of Papuan ancestry may have dispersed into Wallacea, with the primary source being more closely related to some modern West Papuan groups.

### Evidence for retention of AMH founder ancestry in Wallacea

While our ADMIXTURE analyses support multiple sources of Asian- and Papuan-related ancestry being common across Wallacea, evidence for the retention of founder AMH ancestry in Wallacea is less obvious – potentially due to a lack of suitable genetic proxy for this deep ancestry source in this analysis. A hint that deep AMH ancestry is present in modern Wallaceans is provided by two ancestry sources observed at levels <5% in our best-fitting ADMIXTURE models (i.e., 9 ≥ *K* ≥ 11) that reach their highest frequencies in groups from the Andaman Islands (Onge) and Philippines (Aeta Negritoes) (i.e., salmon and red components in **Figs. 2A and S2**) – though ancestries at such low levels may also represent modelling artefacts. Similarly, projection of the recently reported ∼7,000 year old Tolean forager (19) within the full 844 sample PCA resulted in an isolated central position in the first two PCA dimensions (**Figs. 2B and S3**), which is consistent with previous estimates of their ancestry being a mix of an unknown Asian and Near Oceanians lineages (19) but does not provide direct evidence for AMH ancestry in Wallacea.

To explore the presence of founder AMH ancestry in modern Wallacea further, and differentiate this from Melanesian-specific sources (i.e., all Papuan populations and Baining from the Bismarck Archipelago), we used qpAdm (38) to identify plausible ancestors and infer their relative contributions to each Wallacean population. Following results from our ADMIXTURE and PCA analyses, we included multiple Asian and Melanesian populations (limited to a subset of Papuan language speakers; **see SI Methods)** as potential ancestry sources, and added the Toalean forager and Onge population as proxies for founder ancestry. To improve qpAdm sensitivity and power, we rotated populations across ‘left’ and ‘right’ groupings to produce a large set of ancestry models for each Wallacean population that were evaluated for plausibility (**see SI Methods;** (39)).

In keeping with our ADMIXTURE and PCA results, plausible qpAdm models comprising at least one Asian and one Papuan ancestry source were identified for all 14 Wallacean groups (**Fig. 3, Table S2**), with the Indigenous Taiwanese group appearing in all models (range ∼30**–**80%) and, less expectedly, the West Papuan Sorong group being most commonly occuring Papuan source (range ∼10**–**50%) (**Fig. 3B**). The two most common qpAdm models combined these two ancestries with a third source – either the Toalean forager or the Onge (featuring in 9 and 8 Wallcean populations, respectively; **Fig. 3B) –** and these three-source models were among the most parsimonious ancestry configurations for 11 of the 14 Wallacean populations (**Fig. 3B)**. The presence of this third group in plausible qpAdm models for most Wallacean populations is consistent with the widespread retention of AMH founder ancestry across the archipelago, with levels peaking in populations from western Wallacea (i.e., Sulawesi and Flores) and dropping below 5% in groups in the east of the archipelago (**Fig. 3C**). However, further interpretation of the genetic origins of this third ancestry source is complicated by the presence of multiple equally parsimonious models for most Wallacean populations, as well as the lack of phylogenetic context for the Onge- and Toalean forager-related ancestries. Accordingly, we turned to admixture graph modelling to situate the different strands of modern Wallacean ancestry within a robust population genetic framework.

**Figure 3.**
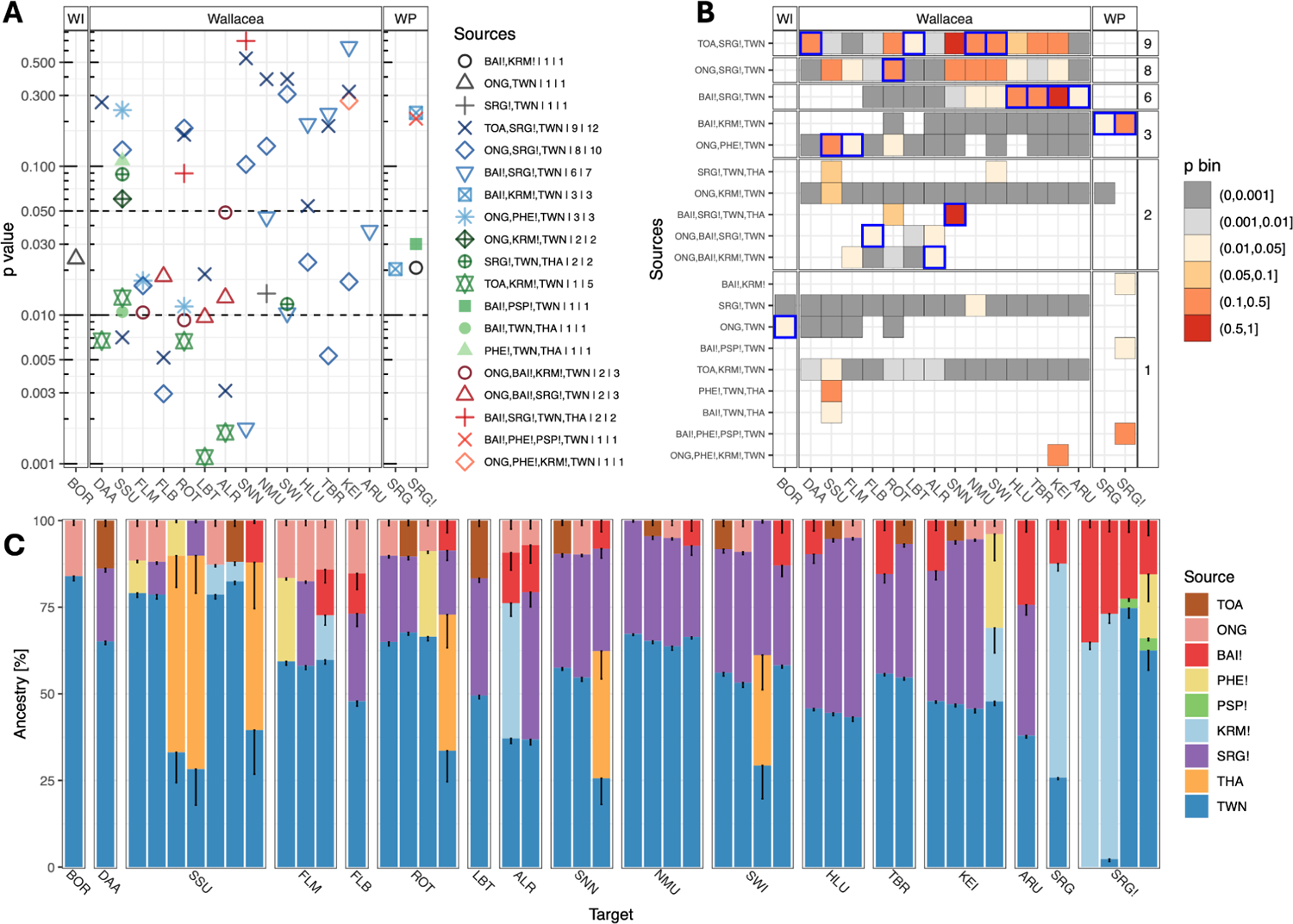
Widespread Papuan and AMH founder ancestry in Wallacea. **A.** Ocean Plot of qpAdm results, showing *p* values for all ancestry models with *p* > 0.01 in at least one of the target populations (shown on x-axis). Inference for qpAdm involves testing whether a specific set of ancestry sources (see key) is *insufficient* to capture the population genetic profile of a target population, whereby models rejecting this hypothesis (*p* > 0.01) are considered as a plausible set of genetic ancestors for the target population (WI: Western Indonesia; WP: West Papua). **B.** Two qpAdm models were identified model for 11 of the 14 Wallacean populations (y axis), suggesting ancestral sources related to the Indigenous Taiwanese and Sorong in West Papua, and either the Andamanese Onge or Toalean foragers were widespread across Wallacea (see key for *p* value bins; blue boxes denote highest *p* values for each population). The exceptions included one group that (Lembata, LBT) was more parsimoniously modelled as a combination of Indigenous Taiwanese and Sorong ancestries, with another (Aru, ARU) additionally including Baining, and the other two (Alor and Flores Bena, ALR and FLB, respectively) requiring a minimum four sources (e.g., including Baining and Onge in addition to Indigenous Taiwanese and Sorong; Fig. 3B).**C.** Fractional ancestry contributed by each source group (y axis) for all plausible models in each target population. Estimation uncertainty (black whiskers) is particularly high for estimates from Thai groups that represent Southeast Asian ancestry in our qpAdm models, suggesting that this population may be a poor genetic proxy of the putative ancestral population.

### Back-migrations from New Guinea spread Papuan ancestry across Wallacea

To develop a parsimonious model of the separations and mixing events underlying Wallacean population genetic history, we used the recently developed *find_graph* algorithm to identify well-fitted admixture graphs from the vast model space (40). Initially, we inferred an admixture graph for all populations identified in our previous analyses as potential contributors to Wallacean ancestry—i.e., Indigenous Taiwanese, Thai, Baining, Onge, Aeta, the Toalean forager, and multiple Papuan populations—with archaic hominin genomes added to capture additional historical mixing events, and a handful other human groups (Han Chinese, Nigerian Mbuti, Aboriginal Australian) included to provide further phylogenetic context. This initial admixture graph served as the topological scaffold to which subsets of Wallacean populations were later appended, also using *find_graph* (**see see SI Methods**).

The best fitting admixture graph scaffold (all absolute standardised *f*_3_ residuals, |*Z|,* less than 3; **Fig. S5 & S6**) identified by *find_graph* reproduces fundamental features from a previously reported graph capturing the global radiation of AMH beyond Africa (41), including the effectively simultaneous split between Asian (combining Indigenous Taiwanese, Thai, and Han), Onge, and ‘Southern’ (combining Aeta, the Toalean forager, and Near Oceanians) lineages early during this dispersal, and the introgression of ∼4% Denisovan ancestry into the Southern lineage that is absent in other AMH groups (**Figure 4, S5 & S6**). We also confirm the recent finding (19) that the Toalean forager derives a large portion of their ancestry (∼35%) from an unknown deeply divergent AMH group, with the remainder coming from a lineage that is genetically equidistant from Aboriginal Australian and Melanesian populations (**Figs. 4, S5 & S6, Table S5, see SI Methods**), supporting claims that the latter represents a founding Wallacean AMH lineage that split from the ancestors of modern Near Oceanians prior to the peopling of Sahul.

**Figure 4.**
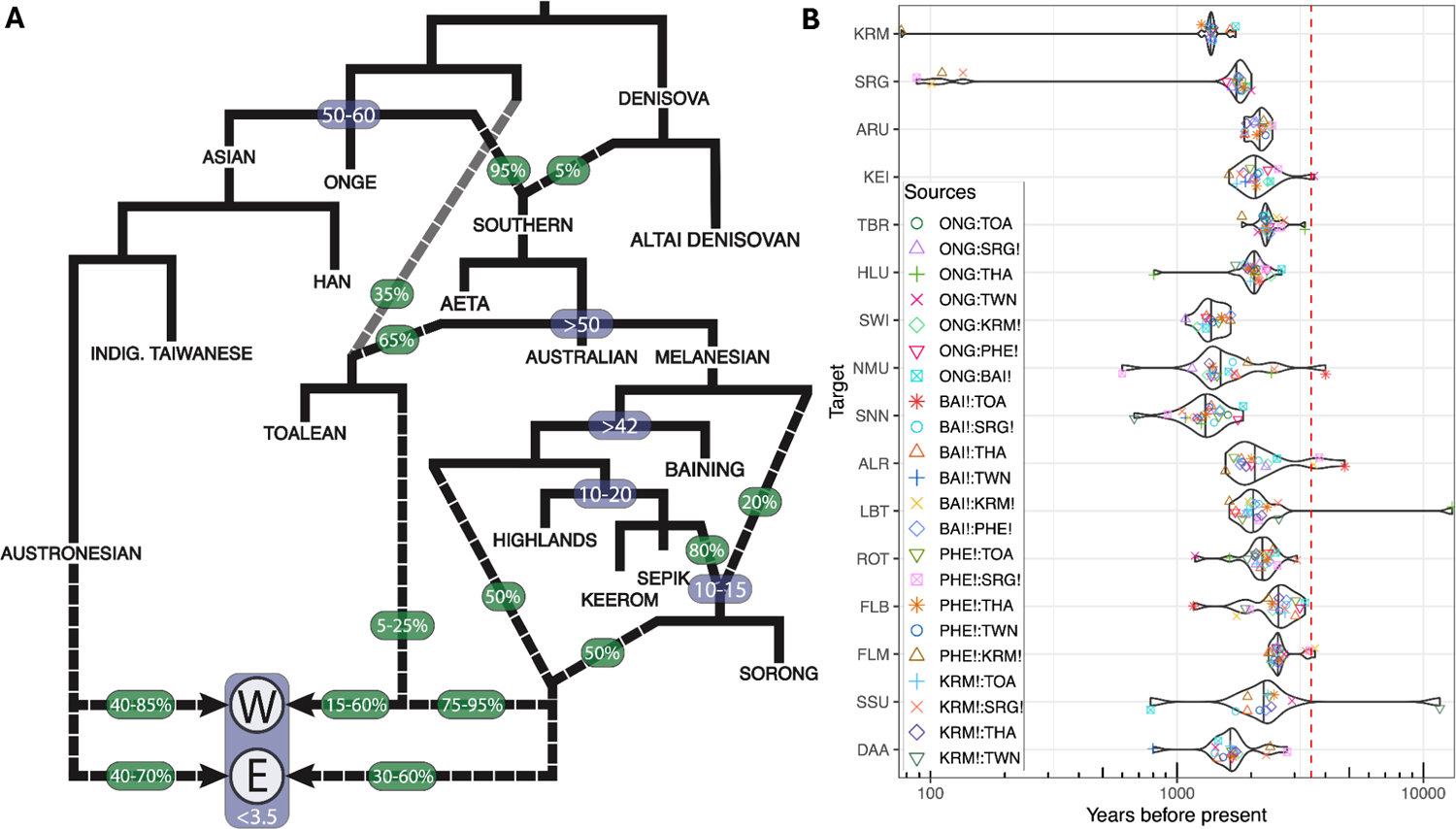
Three genetically distinct Papuan ancestries are widespread across Wallacea. **A.** Summary of well-fitting admixture graphs (all standardised residuals < 3) detailing key separations (solid lines) and mixing events (dashed lines) contributing to Wallacean ancestry (note that branch lengths are not representative of drift relationships). Approximate admixture proportions and dates (kya) for key population separations and mixing events are highlighted in green and blue boxes, respectively. All modern Wallaceans are descendants of lineages related to Austronesian seafarers and three genetically distinct Papuan groups, with groups from western and central Wallacea (W; South Sulawesi [SSU], Daa [DAA], Flores Bena [FLB], Flores Manggarai [FLM], Alor [ALR], Lembata [LBT], and Rote [ROT]) also drawing ancestry from a lineage related to Toalean foragers that is not detected in eastern populations (E). Slightly worse fitting models are obtained when the basal AMH lineage indicated by the transparent line is removed from the ancestry of modern Wallaceans, suggesting it may be specific to Toalean foragers. Note that the Filipino Aeta group also have Austronesian-related ancestry that is omitted here for clarity. **B.** MALDER admixture date estimates using proxies for Wallacean ancestor source groups (see key). Dates suggest that mixing only commenced after the arrival of Austronesian seafarers – with ∼97% of dates falling after 3.5 kya (red line) – though the tight clustering of different pairs of ancestor proxies within populations suggests that admixture was ongoing and initial mixing events may predate this period.

An intriguing result that is not evident in previous studies is the phylogenetic position of an unknown lineage comprising ∼20% of the ancestry of the Papuan language-speaking Sorong population, which splits from other Near Oceanians following the separation of Aboriginal Australians but prior to the diversification of Melanesian populations (**Figs. 4, S5 & S6**), hinting at a possible Wallacean origin if the ancestors of Aboriginal Australians and Papuans arrived in Sahul in separate movements (42–44). However, additional evidence suggests that a New Guinea origin for this lineage is more parsimonious, as the remainder of Papuan language-speaking Sorong ancestry comes from a source that is equally related to both Sepik and Keerom populations, with qpAdm testing confirming this distinctive combination of Melanesian lineages as the predominant ancestry sources in Sorong (**Figs. 3**). This phylogenetic context suggests a scenario where a deeply divergent Papuan group emerged around the Bird’s Head Peninsula in the interval following the peopling of Sahul and prior to the settlement of the Bismarcks by ∼42 kya (45), and subsequently mixed with a migrant group that likely moved eastwards from their homelands in central-northern New Guinea (this group being equidistantly related to the Keerom and Sepik groups in central-north New Guinea; **Figs. 4, S5, & S6, also see Discussion**).

Subsequent fitting of Wallacean populations to our admixture graph scaffold using *find_graph* resulted in a series of well-fitted models (maximum |*Z| <* 3 for all models; **Figs. S7–S14)** that infer a population genetic history consistent with our previous analyses. These admixture graphs reiterate the core findings of our qpGraph results, including the finding that several western and central Wallacean populations derive ancestry from a source related to the Toalean forager, though this is limited to western and central Wallacean populations in our admixture graphs (**Figs. 4, S9, S11 & S13**). Notably, equally well fitted models were obtained when modern Wallaceans were forced to split from the Toalean lineage before it received the deeply-divergent AMH ancestry (maximum *|Z|* < 3; and *p* = 0.52 for bootstrap model fit comparison; **Table S5, see SI Methods**), indicating that this enigmatic AMH group may not contribute to the genetic profile of modern Wallaceans. Additionally, all Wallacean populations share three distinct ancestry sources related to Indigenous Taiwanese (i.e., Austonesian seafarers), West Papuan Sorong, and another unknown early-diverging Near Oceanian lineage (**Figs. 4 & S7–S14**). The phylogenetic position of this unknown lineage—branching from other Melanesian groups after the divergence of Baining but before Mappi separates—suggests an source from New Guinea that diverged from other mainland Papuan groups prior to previously reported internal separations by 10 kya (28, 42).

These results imply that all modern Wallaceans are descendants of at least three genetically distinct Papuan lineages that collectively comprise between 15% to 60% of modern Wallacean ancestry (**Figs. 4 & S7–S14**). Contributions from local AMH founders are comparatively small (< 10% of ancestry of each Wallacean population; **Figs. 4 & S6–S14**) and appear to contribute little to no ancestry to populations located in the east of the archipelago, corroborating the general absence of basal Wallacean lineages in large phylogenies including Australo-Melanesians (18, 22). Accordingly, our results imply that the distinctive Papuan ancestry cline observed in modern Wallacea was largely created by back-migrating Papuan lineages that became less impactful in the eastern margins of the archipelago, and suggest that AMH founder ancestry may in fact be distributed in a countergradient to this dominant Papuan ancestry cline.

### Dating historical admixture events

A feature shared by all of our best-fitting Wallacean admixture graphs is the consistent ordering of the different admixture events, with the two Papuan-related ancestries always mixing prior to admixture with the Indigenous Taiwanese-related lineage, and admixture involving the Toalean forager-related group occurring between these two mixing events (**Figs. 4 & S7-S14**). In some instances these admixture events occur simultaneously—i.e., no intervening drift branches—implying separation by time intervals too brief to generate diagnostic genetic differences. Notably, this simultaneity is restricted to admixture events combining the Toalean forager-related lineage with the two Papuan-related ancestries, with subsequent mixing of Indigenous Taiwanese-related ancestries tending to occur after a period of shared drift. A parsimonious interpretation of these patterns suggests that the two unknown Papuan lineages mixed in New Guinea prior to dispersing across Wallacea where they mixed with local lineages related to the Toalean forager, and that both events preceded the arrival of Austronesian seafarers.

To explore the timing of these historical admixture events in more detail, we applied MALDER (v1.0; (46)) to different pairs of putative ancestors for each Wallacean population (**see Methods**). Strikingly, ∼97% of plausible MALDER estimates occurred after 3.5 kya (**Fig. 4, Table S3**), with admixed West Papuans having similar dates, implying that the modern genetic diversity in both regions arose from admixture events that postdate the arrival of Austronesian seafarers. Because Wallacean populations likely experienced successive bouts of admixture beyond initial mixing events, which can bias estimates toward the more recent events rather than the initial mixing period (47), we re-estimated admixture timings for a subset of Wallacean and West Papuan populations using LaNeta (48), an approach that accommodates two temporally distinct pulses of introgression from one of the proxy ancestors (**see Methods)**. However, while this resulted in multiple pre-Austronesian mixing dates (**Fig. S15, Table S4**), these included several seemingly anachronistic events (i.e., admixture involving Austronesian seafarer proxies prior to 3.5 kya), pointing to potential estimation issues. Accordingly, our admixture timing estimates strongly support historical movements and mixing events in Wallacea involving both Papuan migrants and Austronesian seafarers in the past 3,500 years, though earlier mixing dates remain plausible and further evaluation requires contextualisation with existing archaeological and linguistic records (see next section).

## DISCUSSION

Our survey of 256 newly sequenced genomes from Wallacea and West Papua provides a revised view of Wallacean genetic history, revealing that the characteristic longitudinal cline of Papuan-related ancestry largely results from the back-migration of three genetically distinct lineages from New Guinea, rather than differential descent from AMH founders. The ordering admixture events between the three Papuan ancestries indicate that these predate mixing with local Wallacean populations and Austronesian seafarers, suggesting both mixing events likely occurred in New Guinea prior and the westward dispersal of the resulting admixed lineage into Wallacea. Our LaNeta estimates suggest that the first of these mixing events, which involved the Papuan ancestors of the Sorong group, occurred between 10–15 kya (**Fig. S15**), a period that coincides with the initial genetic separation of populations currently living in highland and lowland regions of PNG (28, 42) and shifting settlement patterns triggered by the submersion of coastal landscapes and improved accessibility to high altitude valley systems due to warming temperatures (49). These findings indicate that the end of the LGM was a period of major demographic change in New Guinea and suggest that the observed cluster of LaNeta mixing dates falling between 10–20 kya in the present study (**Fig. S15**) may capture contemporaneous events in mainland New Guinea rather than attesting to Papuan migrations into Wallacea.

### A West Papuan interaction hub may have facilitated dispersals into Wallacea

Our MALDER admixture time estimates suggest that the historical Papuan migrations into Wallacea occurred in a series of movements that followed the arrival of Austronesian seafarers within the past 3.5 years, echoing findings from earlier population genomic studies (12, 14, 15, 20). These timings closely match the first appearance of the New Guinea endemic cuscus, *Phalanger orientalis*, in Timor around 3.3 kya, likely through human translocation (23), with linguistic analyses indicating that words for ‘cuscus’ reflecting #mansər likely diffused into East Wallacea through a chain of languages that originated in New Guinea (50). The patterning of our MALDER dates—with substantially more date variation between Wallacean populations than within them (**Fig. 4**)—also echo the staggered appearance of Austronesian-associated red-slip pottery in archaeological sites across Wallacea (51) and suggest that Papuan migrants and Austronesian seafarers arrived in each island contemporaneously, though initial arrival times likely differed across islands.

Converging archaeological and linguistic evidence suggest that West Papua once served as a regional hub for cross-cultural interactions between local Papuan groups and Austronesian seafarers, with the Bomberai Peninsula potentially acting as a launching point for historical migrations into Eastern Wallacea. For instance, near-identical rock art motifs appear in the Tutuala region of Timor-Leste and across the Bomberai Gulf region of West Papua that, while undated, fit within the ‘Austronesian Painting Tradition’ that arose in the Late Holocene (24) and recent linguistic work has established a genealogical relationship between Papuan languages spoken in Bomberai Peninsula and across Timor, Alor, and Pantar (27), positing Bomberai as a hub in the inter-island dissemination of Papuan languages, culture, and genes across eastern Wallacea. The longitudinal cline in Wallacean ancestries shows that these historical migrations cannot be reduced to the spread of a single admixed source, however, with current genetic evidence supporting a more complex history involving multiple phases of mixing—including lineages related to Southeast Asians that are poorly resolved in the present study (see next section)—that likely continued into the past 1,000 years.

### Evidence for mixing before and after initial Austronesian contact

The Wallacean archaeological record shows signs of heightened population turnover and mobility in the period postdating the arrival of Austronesian seafarers, with activities peaking during the Historic Period in response to intensified inter-clan conflicts and the emergence of slavery and spice trading networks (52).

These demographic changes likely facilitated opportunities for admixture beyond the initial mixing events in many Wallacean islands, a scenario supported by paleogenomic evidence showing that an as yet unknown group related to Southeast Asians mixed with local groups in Flores by 2.5 kya, before reaching the North Moluccas in the past 1,000 years, in their eastward radiation across the archipelago (20). Our qpAdm analyses also produced several plausible models that include both Thai and an Indigenous Taiwanese sources (**Fig. 3**), agreeing with previous findings that Southeast Asian-related ancestry is a distinct component of the modern Wallacean genetic profile rather than another proxy for Austronesian-related ancestry (12, 20). While the origins of the Southeast Asian-related group remain obscure, the timing of reported mixing events are consistent with the spread of Dong Song drums from mainland Southeast Asia during the Metal Age (53) and substantiate the ongoing dissemination of genetic lineages and cultures across Wallacea throughout the Late Holocene.

An important consequence of ongoing historical admixture in Wallacea is that our estimated admixture timings may be biased towards the present (47), such that pre-Austronesian mixing events remain possible. This scenario gains some credence from our admixture graphs, where mixing between Papuan and local Wallacean ancestries are often followed by a period of drift before the introgression of Austronesian seafarer ancestry, denoting a temporal separation between these two mixing events—though the interceding branch length is not directly translatable into time (**Figs. 4 & S7-S14**). Papuan back-migrations into Wallacea prior to the LGM are not readily supported by current archaeological records, however, with evidence from early AMH occupation sites on large Wallacean islands (dated at 48–44 kya; (54–56) suggesting cultural continuity until the onset of the LGM. Records become sparse during the LGM, though the reduced usage of previously occupied sites may signal the relocation of local residents to track resources along receding coastlines rather than regional depopulation events (57).

The post-LGM period marks the onset of a more dynamic period in eastern Wallacean history, with evidence supporting the emergence of inter-island exchange networks for obsidian (58, 59) and maritime technologies (e.g., fishhooks and ground shell adzes; (60)) that coincide with the initial settlement of small islands and new and intensified site usage on larger islands (57). These networks may have linked eastern Wallaceans to coastal West Papuan groups, as suggested by the presence of shared cooking technologies (23) and the westward spread of banana cultivars from New Guinea (61), and could have resulted in the resettlement of Papuan peoples in newly occupied Wallacean sites. While this would explain the apparent absence of local AMH founder ancestry in modern eastern Wallacean populations, these archaeological patterns are typically interpreted as evidence for the expansion of pre-existing Wallacean groups reacting to shifting coastal landscapes and resource dependencies as sea levels rose throughout the Terminal Pleistocene (11). Accordingly, current evidence supporting the pre-Austronesian movement of Papuan technologies and genes into Wallacea is suggestive rather than substantive and further investigation is needed to resolve this question.

### Wider regional impact of Papuan migrations and future work

Previous population genomic studies suggest that the historical spread of Papuan-related ancestry reached the southern Philippines during the Holocene (62) but did not penetrate the eastern Indonesian islands that were formerly part of the Sunda (12, 14). Our analyses provide further support for these patterns with Papuan-realted ancestry appearing at low proportions (between ∼1–5%) in the Ati group from our ADMIXTURE results (**Fig. 2**), and the sole plausible qpAdm model for the western Indonesian Borneo population lacking a Papuan ancestry source (**Fig. 3**). We were unable to find a plausible qpAdm model for the Ati population, and using *find_graph* to append this population into admixture graph resulted in a similar phylogenetic context to the Aeta group (**Fig. S16**), such that further work is needed to confirm if the Papuan ancestry in contemporary populations in southern Philippines and Wallacea shares a common genetic origin.

Our study provides new insights into the genetic history of Wallacea, indicating that Papuan migrants have had an impact on Wallacea’s modern genetic landscape rivaling the contributions of Austronesian seafarers. These two historical migrations have largely replaced local AMH founder ancestry that, in combination with heightened levels of population mobility throughout Wallacea in the past 3,500 years, pose a serious challenge for reconstructions of the original AMH migrations into Sahul that rely solely on modern genetic data. Other outstanding problems include the need for refined timings for the historical diffusion of Papuan peoples within Wallacea and into neighbouring regions, determining if any of these movements and mixing events predate Austronesian arrival, and obtaining a better understanding of the genetic and cultural interactions involving local Wallacean populations and the different migrant groups. Shedding light on these fundamental questions will require further genomic research, particularly paleogenomic studies utilising human remains that predate Austronesian arrival, and synthesis with existing and emerging linguistic, archaeological, and palaeoecological records.

## METHODS

### Sample collection and ethics

Genomic data was consented from 256 individuals from 12 different populations across Wallacea (see full list and associated metadata in Table S1). Permission to conduct the research was granted by the National Agency for Research and Innovation, under the auspices of the Indonesian State Ministry of Research and Technology. Informed consent for all 252 individuals was obtained for the collection and use of all biological samples during community visits that were overseen by researchers from the Eijkman Institute for Molecular Biology, following the Protection of Human Subjects protocol established by the Eijkman Institute Research Ethics Commission (EIREC). All consenting participants were surveyed for their current residence, familial birthplaces, date and place of birth, and genealogical information for the three to four preceding generations. The study is also approved by The University of Adelaide Human Research Ethics Committee (Ethics approval no. H-2020-211).

### DNA sequencing, processing and alignment

DNA extractions and library preparations for all 256 samples were performed at the Eijkman Institute for Molecular Biology laboratory facilities in Jakarta, Indonesia. Extractions were taken from whole blood samples using the Gentra Puregene Blood Core Kit C (QIAGEN), and libraries prepared using the Nextera DNA Flex Library Preparation Kit (Illumina), both using recommended protocols. After quantifying DNA concentrations for each sample using Qubit dsDNA BR Assay Kit (Thermo Fisher Scientific), all 256 DNA libraries were multiplexed into a single pool and submitted to 150 bp paired-end sequencing across three lanes of an Illumina NovaSeq S4 flowcell.

For all samples, raw sequence reads in the fastq format were processed with fastp to remove adapters and trim poly-G and poly-X tails, with additional trimming of terminal nucleotides (5’ end = 20nt, 3’ end = 5nt) using a mean quality threshold of 20 (63). Trimmed reads were mapped and processed following the protocols used for the human genome diversity panel (HGDP) dataset as outlined in ref. (32). Briefly, trimmed reads were mapped to the human reference genome GRCh38 (hg38) using BWA mem v0.7.17 with the T parameter set to 0 (64). Mapped reads were then collated into a single file using samtools v1.9 (65), which was sorted and duplicate reads marked using biobambam2 (66), and bases recalibrated using baseRecalibrator from the GATK software suite v3.5 (67). Variant discovery was performed using GATK Haplotypecaller with parameters “--ERC GVCF’’ and “--includeNonVariantSites’’ to ensure that monomorphic sites were retained and intermediate gVCFs were generated for individual samples. GATK CombineGVCFs was used to amalgamate all gVCFs on a population basis, with joint variant calling on each amalgamated gVCF being performed using GenotypeGVCFs, thereby allowing variant information from all samples in each population to be used in individual genotype calls. This step produced VCF files with phred-scaled likelihoods (PL) —a normalised form of genotype likelihood— which were used for the subsequent imputation process.

### Imputation and genomic dataset merging

To generate a set of robust genotype calls in our low coverage Wallacean and West Papuan genomes, we used the 1000 Genomes Project (TGP) as a reference panel to impute genetic variants using GLIMPSE (31).

Unlike other imputation software, GLIMPSE is designed to work with low coverage genomic data and can improve imputation quality by leveraging genetic information from the target samples, which becomes more advantageous with larger sample sizes (31). The complete set of 252 samples was imputed using GLIMPSEv1.1 (31) with the set of 3,202 phased genomes from the 1000 Genomes Project (TGP) sample collection maintained at the New York Genome Centre (68) as the reference panel. Imputation was performed following the protocol outlined on the official GLIMPSE github repository website using default parameterisations (https://odelaneau.github.io/). First, genomic ‘chunks’ were defined by running the *GLIMPSE_chunk* algorithm on the 3,202 phased TGP genomes that were used as the imputation reference panel, with each chromosome being run independently to speed up computation. Genotype imputation was performed for each resulting chromosome chunk using *GLIMPSE_phase*, with the resulting VCF files for each chunk were then merged using *Glimpse_ligate*, with all imputed loci having genotype probabilities lower than 0.9 sets to missing using the BCFtools plugin (69).

Our low coverage imputed genomes were merged with a global genomic dataset that included publicly available genomes from the Simons Genome Diversity Project (SGDP; (34)) and modern populations from East and Southeast Asia, ISEA, and Near Oceania (see **Table S1**). Merging was performed from VCF files using bcftools v1.9 (69), with all published genomes converted from GRCh37 to GRCh38 coordinates using the liftover tool implemented in Genozip v.12.0.34 ((70), https://genozip.readthedocs.io/dvcf.html), chain file obtained from the UCSC Genome Browser (71)). To this combined set of 855 modern genomes, we added the genome of an ancient Tolean forager from Leang Paninge in Sulawesi (19) (reprocessed the raw sequence data using Eager 2.4.5 (72)), two high coverage genomes from the Denisova and the Altai Neanderthal specimens (53, 54), and the Chimpanzee reference genome (PanTro6; REF), with merging performed by EIGENSTRAT v7.1.2 (73, 74). This final set of 859 individuals was filtered for SNPs missing in more than 5% of the combined samples, or having a minor allele frequency less than 1% across all samples, using PLINK v.1.987; (75), resulting in a set of 3.77M SNPs available for further analysis. Further details regarding the data merging and imputation procedure – and evaluation of imputed genotypes for several of the analytical methods used here – is provided in ref. (33).

### Population genetic analyses

Principal component and ADMIXTURE analyses were performed on the global dataset of 855 modern genomes using a subset of 238,615 unlinked SNPs, with variant pruning performed following Laziridis et al. (76) (**see SI Methods**). Principal component analyses were performed using smartsnp (v1.1.0; (36)), an R package that replicates PCA results from the widely used EIGENSTRAT software (74). Maximum likelihood estimation of discrete ancestry components was performed using ADMIXTURE v1.30 (37), with twenty randomly seeded runs being performed for components ranging from *K* = 3 to *K* = 15. Model fit was determined using the minimum mean cross-validation error measured across 20 replicates (**Fig. SX**). The result for each level of *K* was summarised using Pong (77) with replicates having an average pairwise similarity threshold > 0.9 being treated as supporting the same model. The most frequent model is reported for each level of *K* (**Figs. 2A & SX**). Admixture graph and qpAdm analyses that leverage *f* statistic estimates were performed using the R package ADMIXTOOLS2 (v2.0.0; (78)). Full details of the procedures employed for in our qpAdm and admixture graph inference are reported in **SI Methods**.

### Admixture date estimation

The timing of historical admixture events were estimated using MALDER (46), which measures the decay rate of linkage disequilibrium amongst loci in target populations weighted by the allele frequency divergence in two proxy ancestor populations (79). Target groups included all 14 Wallacean populations and two Austronesian-speaking subgroups of two West Papuan populations. Source populations included Onge and several Asian and Melanesian groups (i.e., Indigenous Taiwanese, Thai, Toalean, Baining, Eastern Highlands, Keerom, Sorong), that were suggested as suitable proxy populations by our population genomic analyses.

Further estimates were obtained by running the recently released software LaNeta, which allows for two pulses of introgression from one of the admixing groups. We limited our analyses to the four Wallacean populations that had the fewest plausible qpAdm models and used the Indigenous Taiwanese, Baining, Sorong, and Onge as test populations. All pairwise combinations were evaluated for each Wallacean population, and we rotated the group contributing two separate pulses of ancestry, resulting in XX models overall. For MALDER estimates, we only included results with *p* values > 0.05 and that had lower 99% confidence bounds greater than 0. Similarly, we only included LaNeta estimates where the first mixing event had lower 99% confidence bounds greater than 0, given that our main focus was in detecting dates predating Austronesian seafarer arrival.

