## Supplementary material for "The genetic origins and impacts of historical Papuan migrations into Wallacea": Figures S1 - S16

**Supplementary Methods**

**qpAdm modelling**

Inference of plausible ancestry models for all 14 Wallacean populations and both Papuan- and Austronesian-speaking subgroups of the West Papuan Sorong population was performed using the qpAdm implementation from ADMIXTOOLS2 (Maier and Patterson 2024). To improve identification of plausible models, the rotation procedure was employed for all qpAdm analyses (Maier et al. 2022; Harney et al. 2021), which explores an exhaustive model test space by cycling potential source populations from one set (i.e. the 'left' cohort) into a second set of populations (i.e. the 'right' cohort). For the Wallacean populations, the left population cohort included Melanesian groups low levels of Austronesian-related ancestry – i.e. Baining, Eastern Highlands, and the Papuan language speaking Keerom and Sorong subgroups – with Onge and the Toalean forager included to capture putative AMH founder ancestry, and Thai and Indigenous Taiwanese as Austronesian seafarer ancestry proxies. The same source populations were used as the left cohort for both Sorong linguistic subgroups, with the exception of Papuan-speaking Sorong and the Toalean forager and addition Mappi and Sepik populations. For all target populations, the right population cohort comprised African Mbuti, East Asian Han, Western Eurasian Sardinians, Indigenous South American Mixe, and archaic Denisovan. This resulted in 501 tests for each Wallacean population, and 256 tests for the two Sorong linguistic subgroups and Keerom with Austronesian linguistic groups. For each target population, plausible models were characterised as those having $p > 0.01$, and estimated ancestry proportions between 0 and 1 (Bergström et al. 2022; Yüncü et al. 2023). Additionally, we removed models where the standard error of the ancestry proportion of a particular source population ($p_{se}$) significantly outweighed the point estimates, $p$ (specifically, excluding models where $p$ falls outside of the 99th percentile confidence interval (i.e. $Z_{99} * p_{se} \leq p \leq Z_{99} * p_{se}$), as such contributions are not statistically discernible from 0.

**Admixture graph modelling**

Until recently, the combinatorial complexity of admixture graph fitting has meant that analyses involving more than a moderate number of populations generally required some degree of manual fitting, with choices made by researchers often leading to suboptimal graphs being identified (Maier et al. 2023). The recently published graph search algorithm, *find_graph* (part of the ADMIXTOOL2 R package; (Maier and Patterson 2024)), offers an automated version of the widely used qpGraph tool that greatly expands the exploration of the vast graph space, leading to improved identification of the best fitting models for complex population histories and mitigating the risks associated with manual approaches (Maier et al. 2023). Accordingly, we used the *find_graph* algorithm to identify well-fitted admixture graphs capturing the genetic history of Wallacean and West Papuan populations.

Despite the computational advantages of *find_graph* over manual fitting, exploration of graph spaces for large and complex population histories is still impractical, such that we split our inference procedure into multiple interlinked steps. The first step involved modelling the genetic history of the three West Papuan populations – i.e. Sorong, Keerom and Mappi, with the first two populations further separated into linguistic subgroups. The model space included a global set of populations informed by our previous analyses, and also included historical admixture events involving archaic Neandertal and Denisovan lineages, rooting the history using a Chimpanzee genome (REF). We ran a *find_graph* on ADMIXTOOLS2 for 19 populations with specific constraints admixture events, including the number (maximum of 15) and ordering (i.e. Altai Neanderthal separates from Denisova prior to the split between Aboriginal Australians and Baining) of events. The best fitting model had a maximum standardised residual, $|Z|$, of 2.97 (**Figs. S5-S11**) (based on $f_3$ statistics of the form $f_3$(population 1, population 2; Chimp)), indicating that it provided a reasonable historical framework for this set of populations.

This Australo-Melanesian admixture graph was used as a scaffold upon which subsets of Wallacean populations were appended, with the admixed Aeta population removed to simplify the model space. Separate *find_graph* procedures were run for the four subsets of Wallacean populations, defined by broad geographical location, namely:

1. Southeast Wallacea, comprising Aru, Kei, and Tanimbar
2. Northeast Wallacea, comprising North Maluku, Sanana, Huaulu, and Sawai
3. South-central Wallacea, comprising Alor, Lembata, Rote
4. West Wallacea, comprising Flores Manggarai, Flores Bena, Daa, and South Sulawesi

Inference involves the same admixture constraints with the West Papuan admixture graph set as the model seed and the Wallacean population subset positioned as a randomly configured clade that separates from the Southern lineage after the Denisovan introgression event. Additionally, based on results from qpAdm models, the Wallacean populations were each constrained to have between one to four admixture events. Importantly, for all subsets of Wallacean populations, the cladal relationships among the population subset were not preserved in the best fitting graph, suggesting that this did not influence the model exploration. The best fitting admixture models identified for some Wallacean population subsets had one or more residuals greater than 3, such that manually modified versions were explored for improved fits, typically involving changing the phylogenetic position of specific lineages that resulted in unidentifiable edges or that were associated with large residuals, or altering the ordering of lineage pairs separated by branch lengths with drift lengths close to

0. In each case, a handful of these changes produced models where all standardised residuals were less than 3, at which point no further modifications were made.

To investigate the robustness of the genetic relationships involving populations from each Wallacean grouping, we also took one population for each Wallacean group (i.e., Aru, Huaulu, Alor, and Daa) and reran the *find_graph* procedure defined above. This resulted in a model (**Fig. S13**) that retained the same ancestry contributions for each Wallacean group that were predicted in other well-fitted models (**Figs. S7-S12**). Our results suggest that the predicted Melanesian and Austronesian ancestry features are shared across modern Wallaceans – differing in their proportional contribution but not in genetic composition – with an additional ancestry source shared with the Toalean forager being common among groups in western and central regions of the Archipelago.

Finally, our ADMIXTURE results suggest that the Ati population had some Papuan ancestry which might be related to the movements detected in previous study (Larena et al. 2021). To investigate this relationship, we appended the Ati group to the Australo-Melanesian admixture graph in the same basal position as used for the Wallacean groups, and reran *find_graph* applying the same model constraints described previously. This produced a well-fitted model (all standardised residuals < 3) where Ati shares the same ancestors as the Aeta, albeit in different proportions. While we do not detect the Papuan ancestry, Filipino populations have a complex population history (Larena et al. 2021) that is poorly represented in our admixture graph analyses that focus instead on Wallacean ancestry, and robustly modelling Ati history requires further work that is beyond the scope of the present study.

**Admixture graph hypothesis testing**

We also used our graphs to test certain hypotheses, which involved changing a specific feature of the graph and comparing the fit of the revised model with the original unaltered version identified by *find_graph*. Specific tests included examining the ordering of separations between the AMH founding groups in Wallacea and Sahul, which have been shown to be equidistant in previous work (Oliveira et al. 2022). To conduct this test, we revised our base Austro-Melanesian graph (**Fig. S5**) by forcing the Melanesian lineage to separate from the common ancestor of Aboriginal Australians and the Toalean forager, with these two lineages then separating and all other features of the original model being retained (**Fig. S6**). This revised model resulted in highly similar residuals (**Fig. S6**) and the two models were statistically indistinguishable when the *compare_fits* function in ADMIXTOOLS2 was used to perform a bootstrap model fit comparison ($p = 0.52$ from 100 bootstrap iterations; **Table S5**). This indicates an effectively simultaneous split among the founding

AMH migrants to Wallacea and the ancestors of Australo-Melanesians that subsequently settled Sahul. Additionally, we tested whether the unknown AMH ancestry source found in the Toalean forager was also present in Wallacean populations that shared ancestry with the former group. This was achieved by modifying the graph shown in Fig. S13, where the deep AMH lineage was forced to introgress into the Toalean forager lineage after it had split from other Wallacean groups (i.e. removing the deep AMH lineage as an ancestor of modern Wallaceans **Fig. S14**). This also resulted in highly consistent residuals (**Fig. S14**) and a non-significant bootstrap model fit comparison ($p$ = 0.42 from 100 bootstrap iterations; **Table S5**), implying that this unknown lineage was not an ancestor of modern Wallacean groups.

### Supplementary Figures

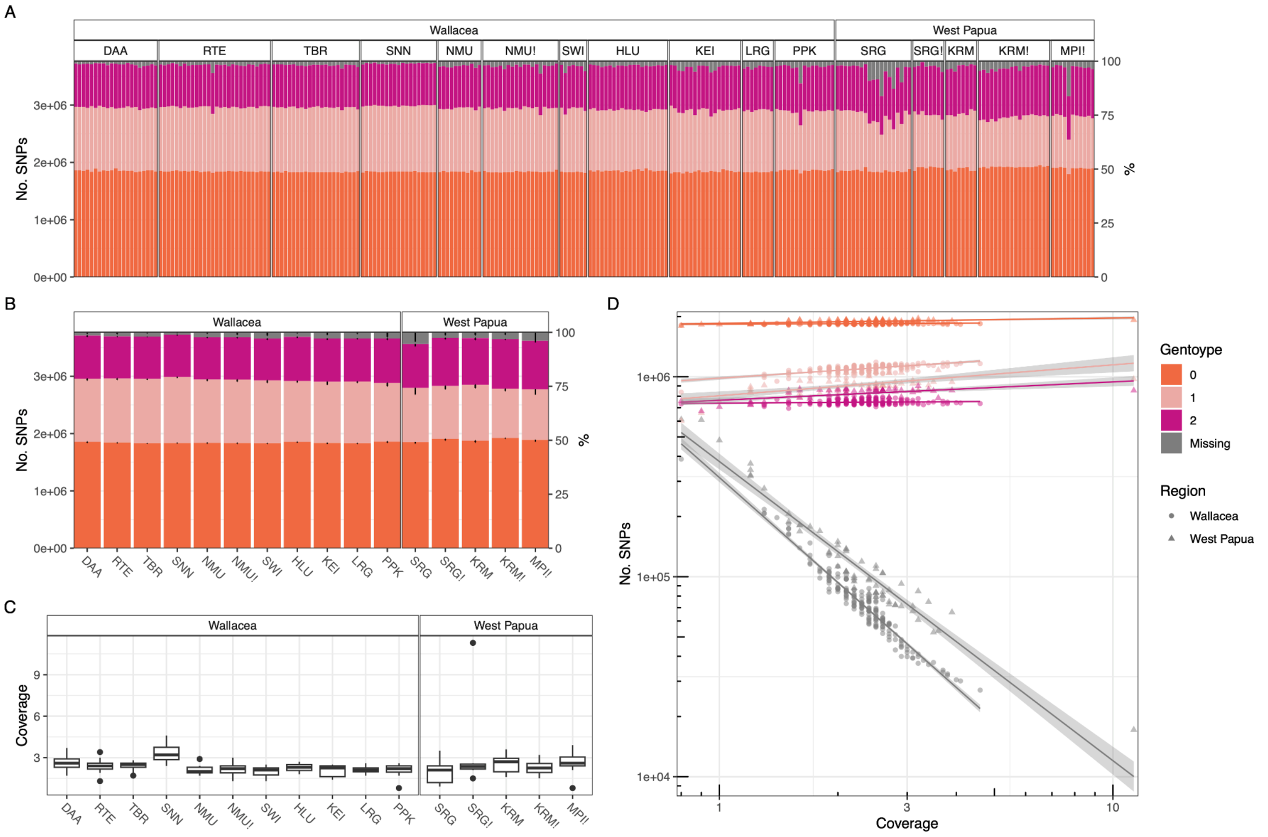


**Figure S1**. **Summary of SNP calling and alignment coverage. A.** Genotype counts for all 276 newly reported Wallacean and West Papuan genomes. **B.** Mean genotype counts and associated standard errors (black bars) for each population. **C.** Box plots showing distribution of genome coverages for each population. **D.** Relationship of genotype genotype counts against coverage, which are approximately linear on the log-log scale. Missingness is inversely proportional to coverage, with West Papuans having more missing SNPs than Wallaceans when fixing the coverage level, with the opposite pattern observed for heterozygote and homozygous alternate allele calls. These results are likely attributable to poorer representation of West Papuans in the imputation panel.

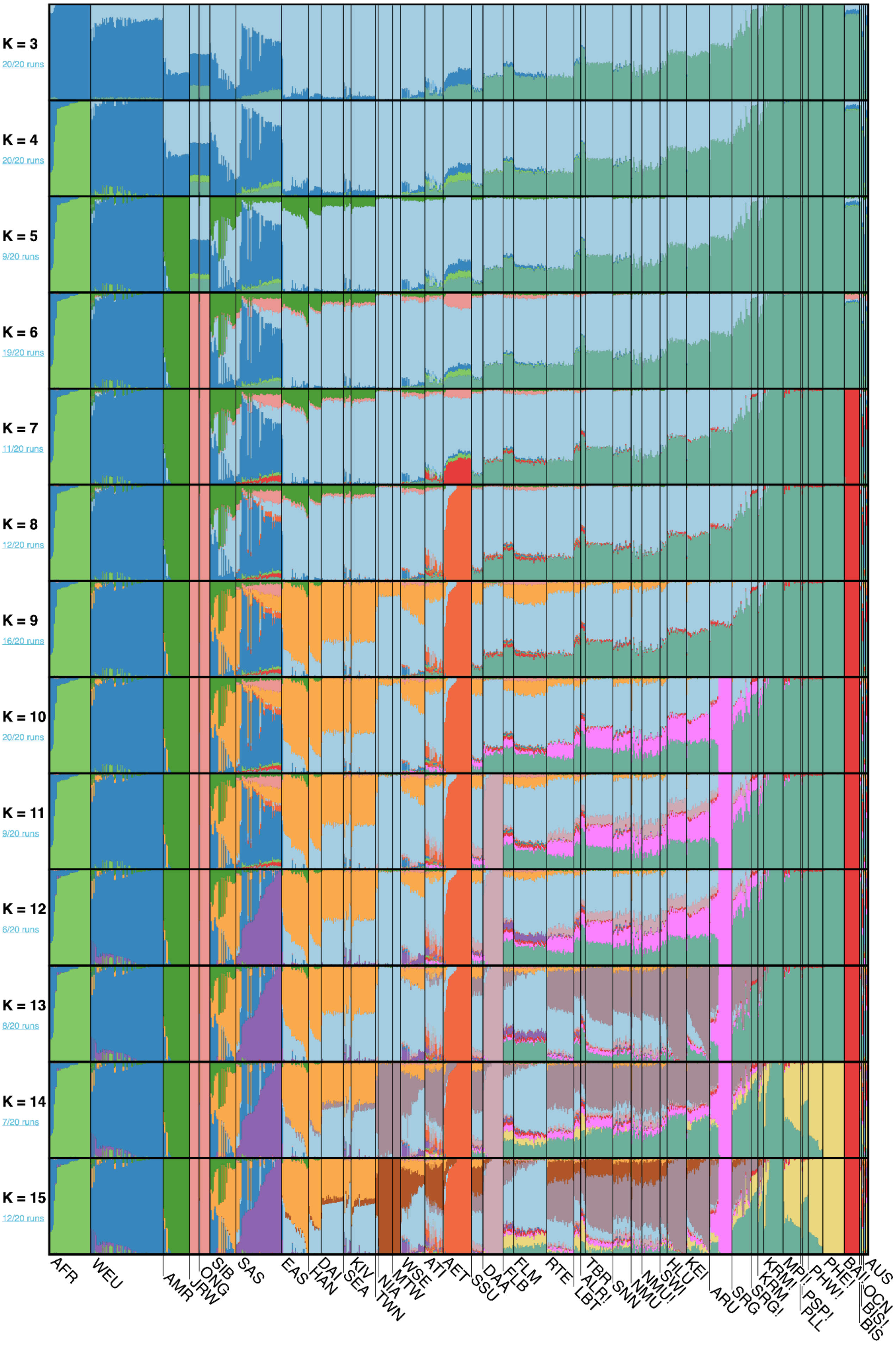

K = 3
20/20 runs
K = 4
20/20 runs
K = 5
9/20 runs
K = 6
19/20 runs
K = 7
11/20 runs
K = 8
12/20 runs
K = 9
16/20 runs
K = 10
20/20 runs
K = 11
9/20 runs
K = 12
6/20 runs
K = 13
8/20 runs
K = 14
7/20 runs
K = 15
12/20 runs
AFR
WEU
AMR
JRW
ONG
SIB
SAS
EAS
HAN
DAI
SEA
KIV
NIA
TWN
MTW
WSE
ATI
AET
SSU
DAA
FLB
FLM
RTE
LBT
ALR!
TBR
SNN
NMU
NMU!
SWI
HLU
KEI
ARU
SRG
SRG!
KRM
KRM!
MPI
PLL
PSP!
PHW!
PHE!
BAI
BIS
BIS!
OCN
AUS

**Figure S2**. **PONG visualisation of ADMIXTURE results.** Ancestry components inferred for varying numbers of ancestral populations, *K*, shown on the left hand side of the plot. Twenty ADMIXTURE replicates were performed for each *K*, with most an example of the most frequently occuring model being visualised in each case (the number of replicates where model was found shown in blue text on left hand side). Different ancestries are shown as different colours, with abbreviated population names shown at the bottom of the plot. Additional details are in **Fig. 2** and **Table S1**. The optimal model occurred at *K* = 9 ancestry components.

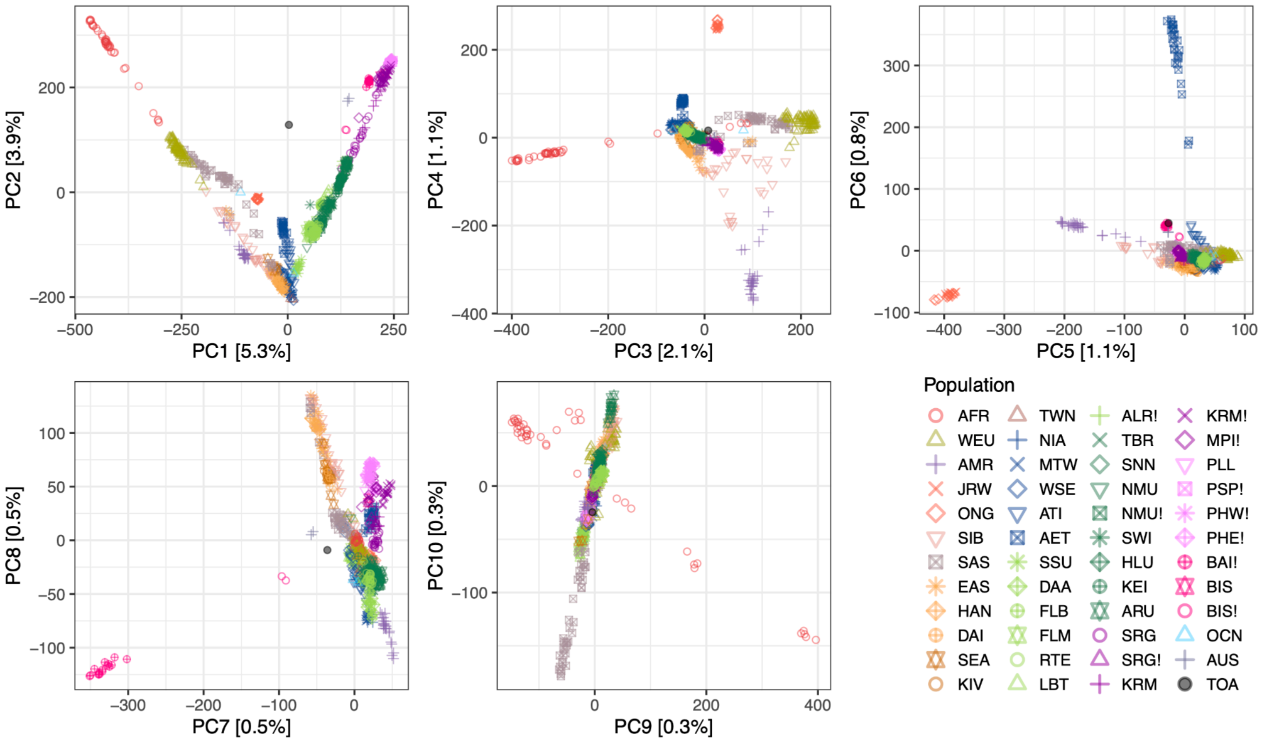


**Figure S3**. **PCA plot of Wallacean and West Papuan genetic diversity contextualised in a global panel.** The first 10 dimensions are highlighted, with the proportion of variance explained in each dimension also indicated in the axes.

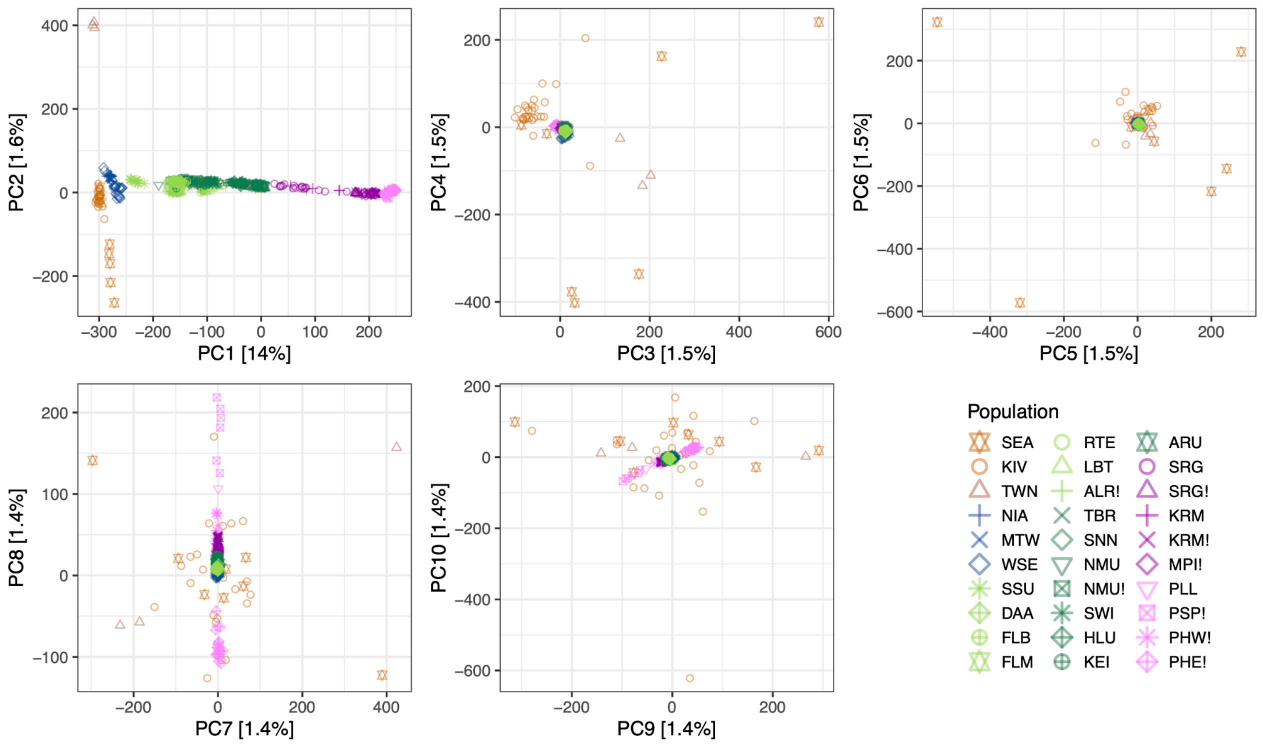


**Figure S4. Projection of Wallacean and West Papuan genetic diversity within PCA built from informative Asian and Papuan groups.** The first 10 dimensions are highlighted, with the proportion of variance explained in each dimension also indicated in the axes. The clustering of Wallacean and West Papuan populations in dimensions beyond the first two indicate that the Asian and Papuan variation captured in these dimensions is not not relevant to differences among the projected genomes.

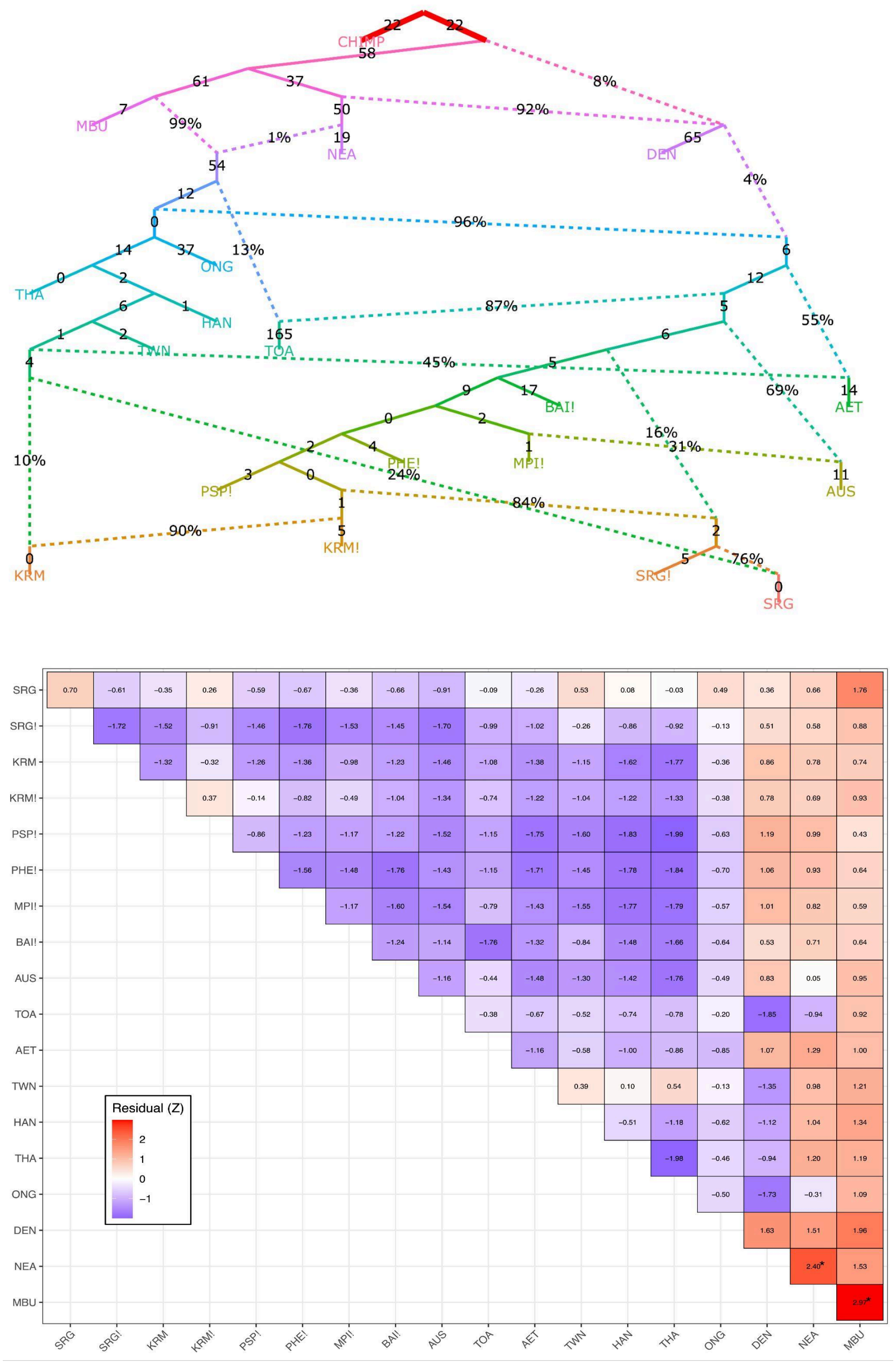


**Figure S5. Admixture graph showing Australo-Melanesian groups within a broad hominin context. A.** Well fitted qpGraph identified by *find_graph* algorithm with minor modifications. **B.** All f3 residuals, |Z|, are less than 3, indicative of a well-fitted graph.

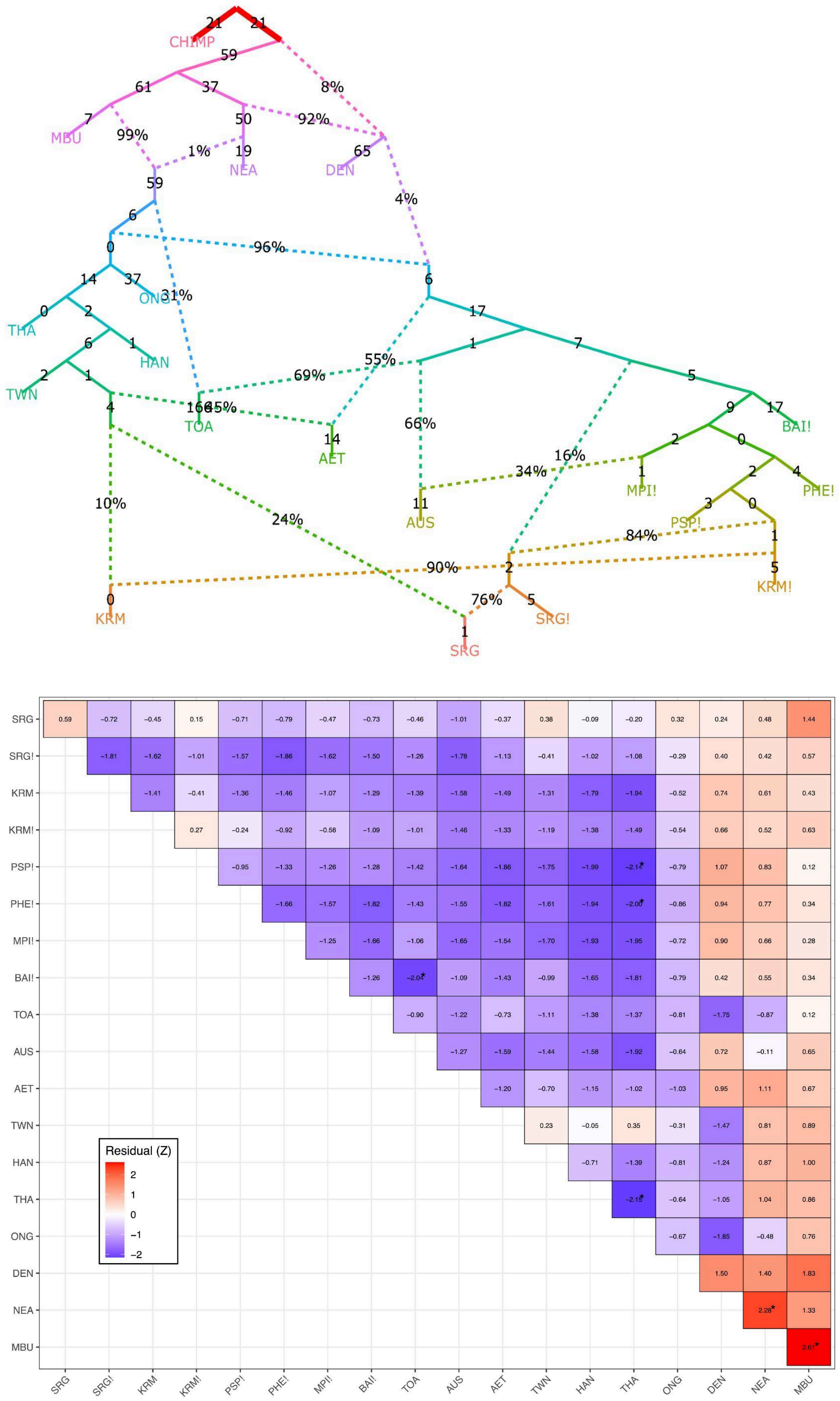


**Figure S6. Admixture graph showing Aboriginal Australians and Toalean foragers having shared ancestry that is absent from Melanesian groups. A.** Same admixture graph as shown in Fig. S5, but changing the ordering of the initial separation events among the founding AMH groups to Wallacea and Sahul. Models in Figs. S5 and S6 are statistically equivalent (see Table S5) indicating equidistant genetic relationships among these populations. **B.** All f3 residuals, |Z|, are less than 3, indicative of a well-fitted graph.

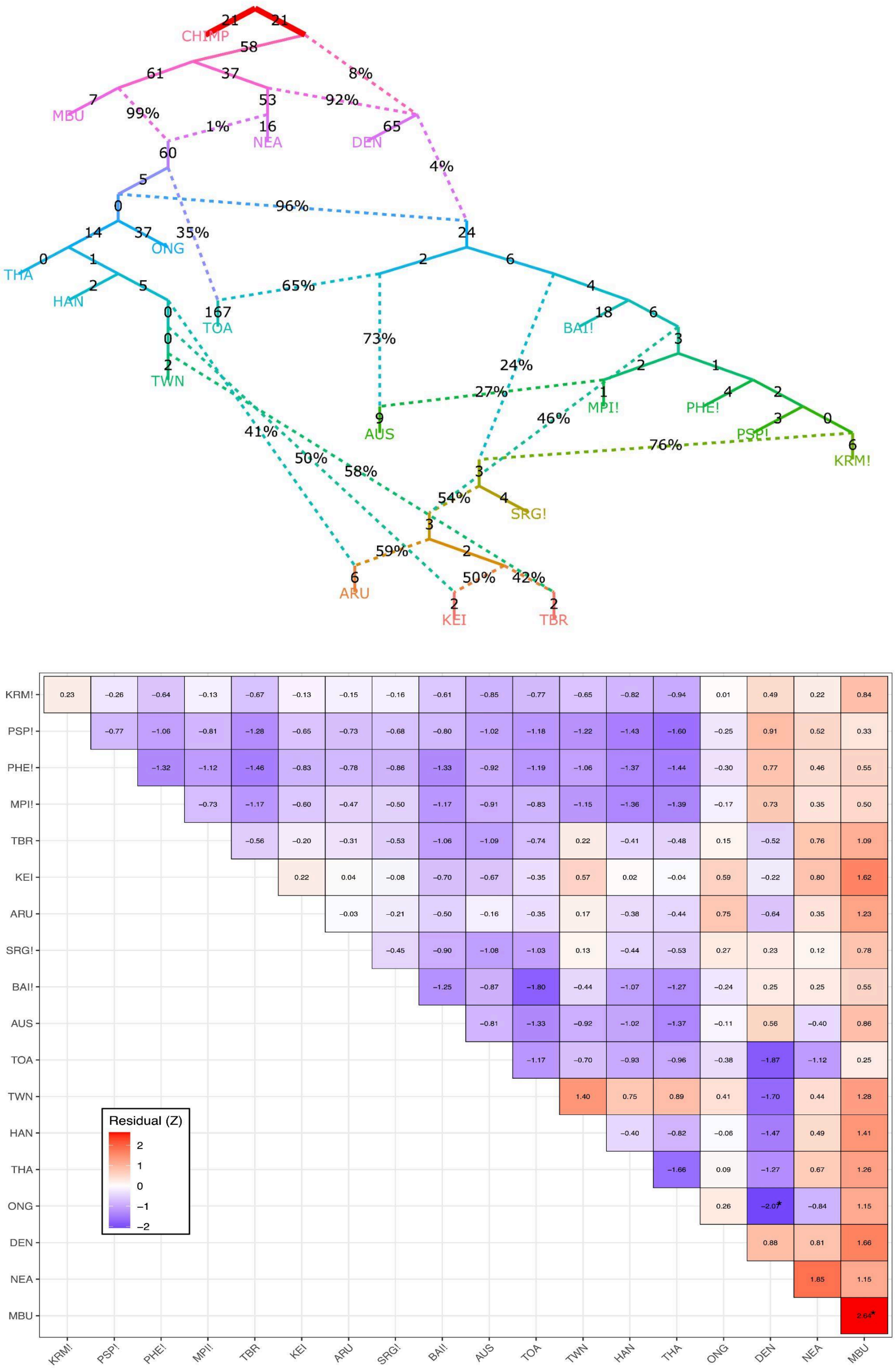


**Figure S7. Admixture graph showing southeast Wallacean groups within a broad hominin context. A.** Well fitted qpGraph identified by *find_graph* algorithm with minor modifications, which includes Wallacean groups from Aru (ARU), Kei (KEI), and Tanimbar (TBR). **B.** All f3 residuals, |Z|, are less than 3, indicative of a well-fitted graph.

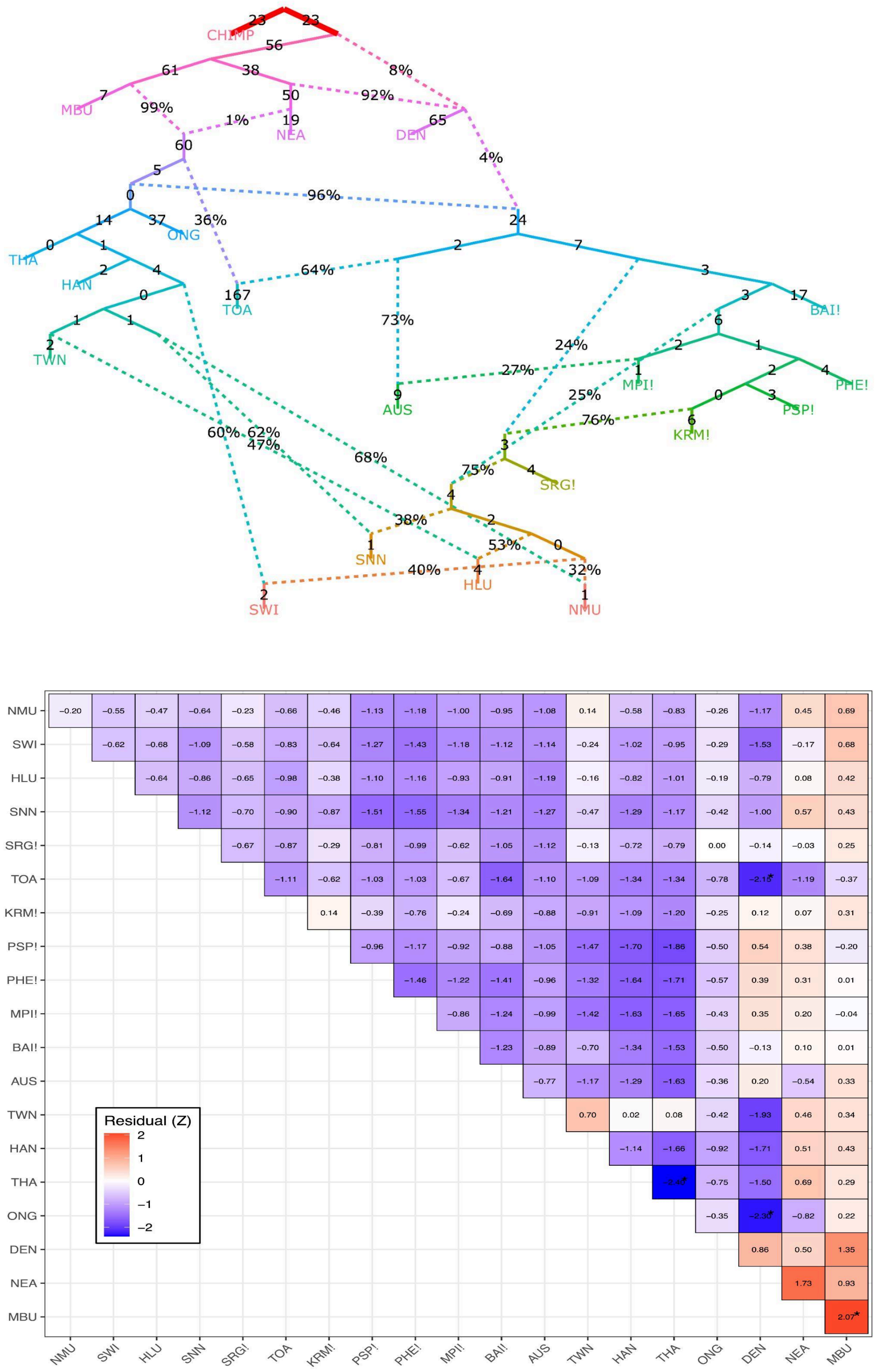


**Figure S8. Admixture graph showing northeast Wallacean groups within a broad hominin context. A.** Well fitted qpGraph identified by *find_graph* algorithm with minor modifications, which includes groups from the North Moluccas (NMU), Sawai (SWI), Huaulu (HLU), and Sanana (SNN). **B.** All f3 residuals, |Z|, are less than 3, indicative of a well-fitted graph.

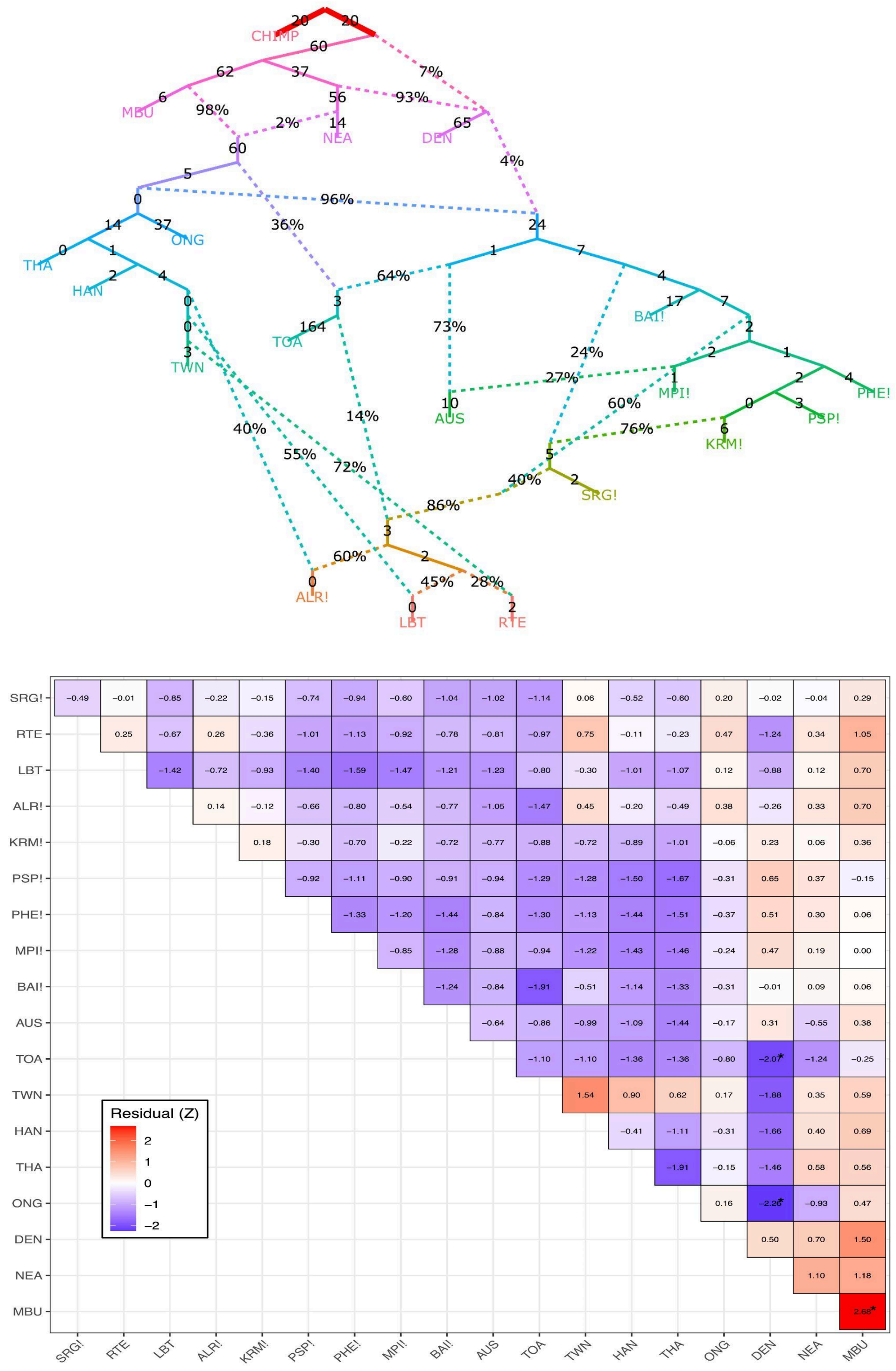


**Figure S9. Admixture graph showing south-central Wallacean groups within a broad hominin context. A.** Well fitted qpGraph identified by *find_graph* algorithm with minor modifications, which includes Wallacean groups from Alor (ALR), Lembata (LBT), and Rote (RTE), and Sanana (SNN). **B.** All f3 residuals, |Z|, are less than 3, indicative of a well-fitted graph.

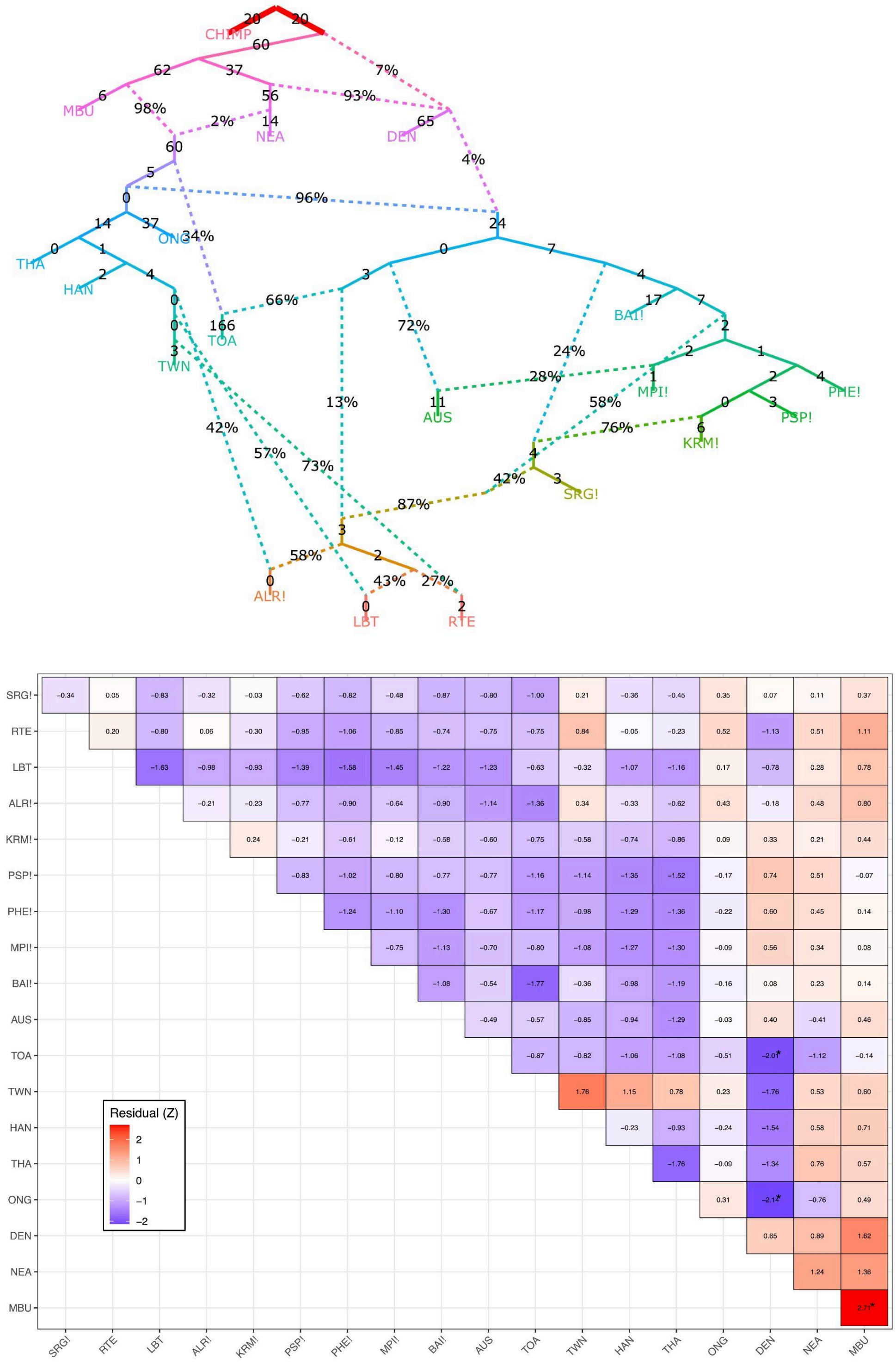


**Figure S10. Admixture graph showing south-central Wallacean groups within a broad hominin context. A.** Well fitted qpGraph identified by *find_graph* algorithm with minor modifications, which includes Wallacean groups from Alor (ALR), Lembata (LBT), and Rote (RTE), and Sanana (SNN) were forced to split from the Toalean lineage before it received the deeply-divergent AMH. **B.** All f3 residuals, |Z|, are less than 3, indicative of a well-fitted graph.

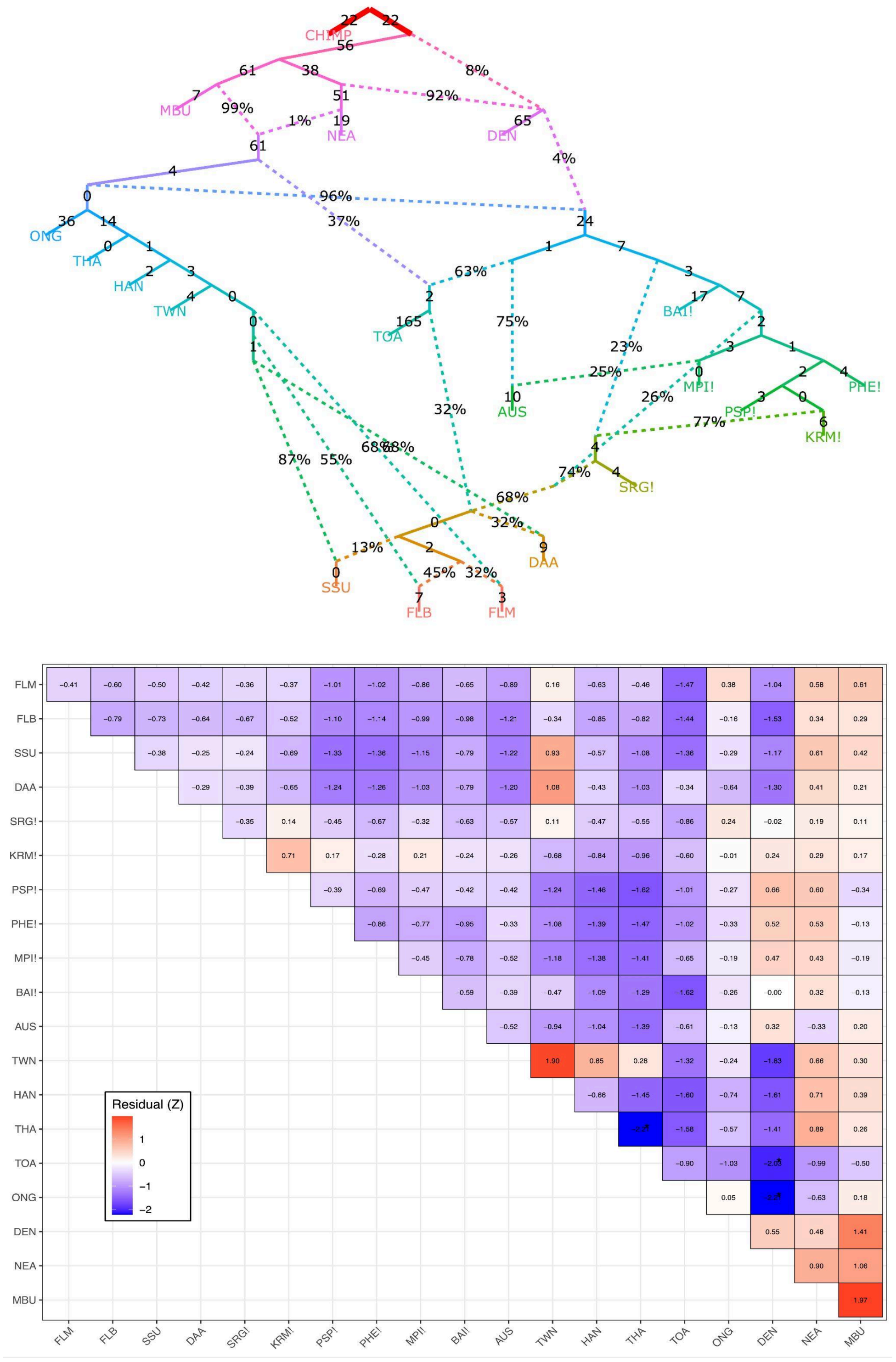


**Figure S11. Admixture graph showing western Wallacean groups within a broad hominin context. A.** Well fitted qpGraph identified by *find_graph* algorithm with minor modifications, which includes Wallacean groups from South Sulawesi (SSU), Daa (DAA), Flores Bena (Flores Bena), and Flores Manggarai (FLM) . **B.** All f3 residuals, |Z|, are less than 3, indicative of a well-fitted graph.

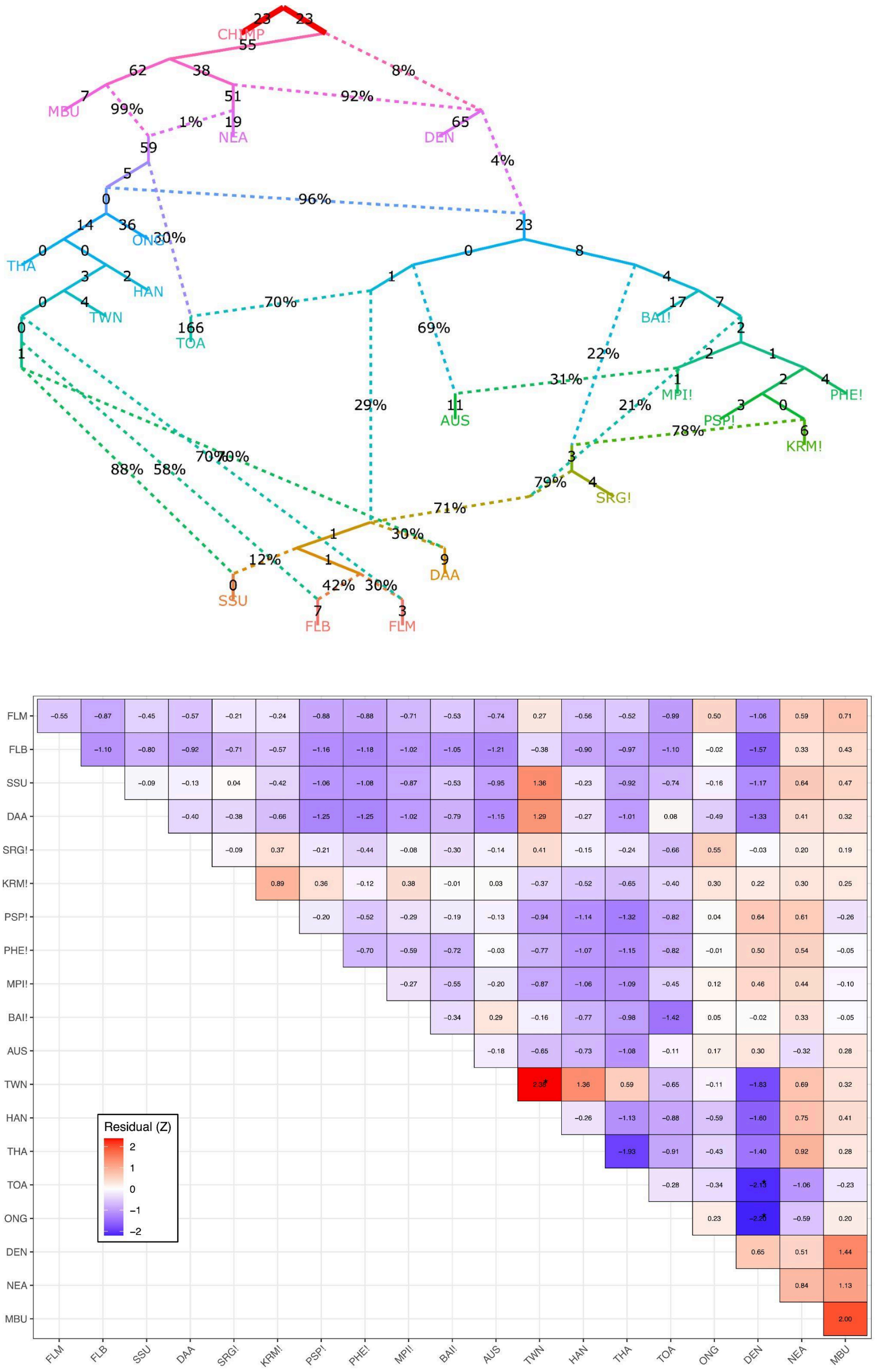


**Figure S12. Admixture graph showing western Wallacean groups within a broad hominin context. A.** Well fitted qpGraph identified by *find_graph* algorithm with minor modifications, which includes Wallacean groups from South Sulawesi (SSU), Daa (DAA), Flores Bena (Flores Bena), and Flores Manggarai (FLM) were forced to split from the Toalean lineage before it received the deeply-divergent AMH. **B.** All f3 residuals, |Z|, are less than 3, indicative of a well-fitted graph.

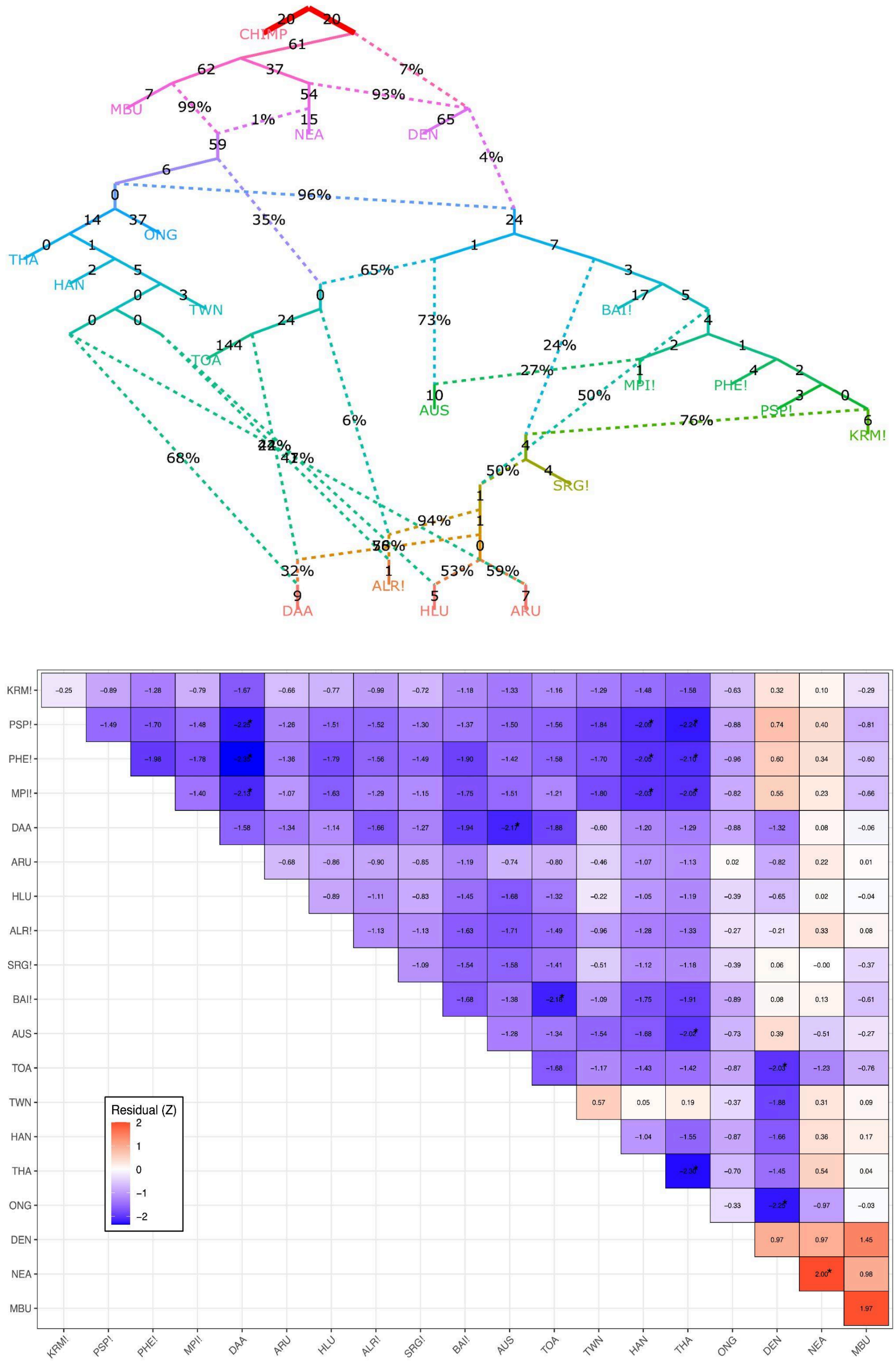


**Figure S13. Admixture graph showing Wallacean groups from each of four geographical subregions. A.** Well fitted qpGraph identified by *find_graph* algorithm with minor modifications, which includes Wallacean groups from Daa (DAA; western Wallacea), Alor (ALR; south-central Wallacea), and Huaulu (HLU; northwest Wallacea), and Aru (ARU; southwest Wallacea). **B.** All f3 residuals, |Z|, are less than 3, indicative of a well-fitted graph.

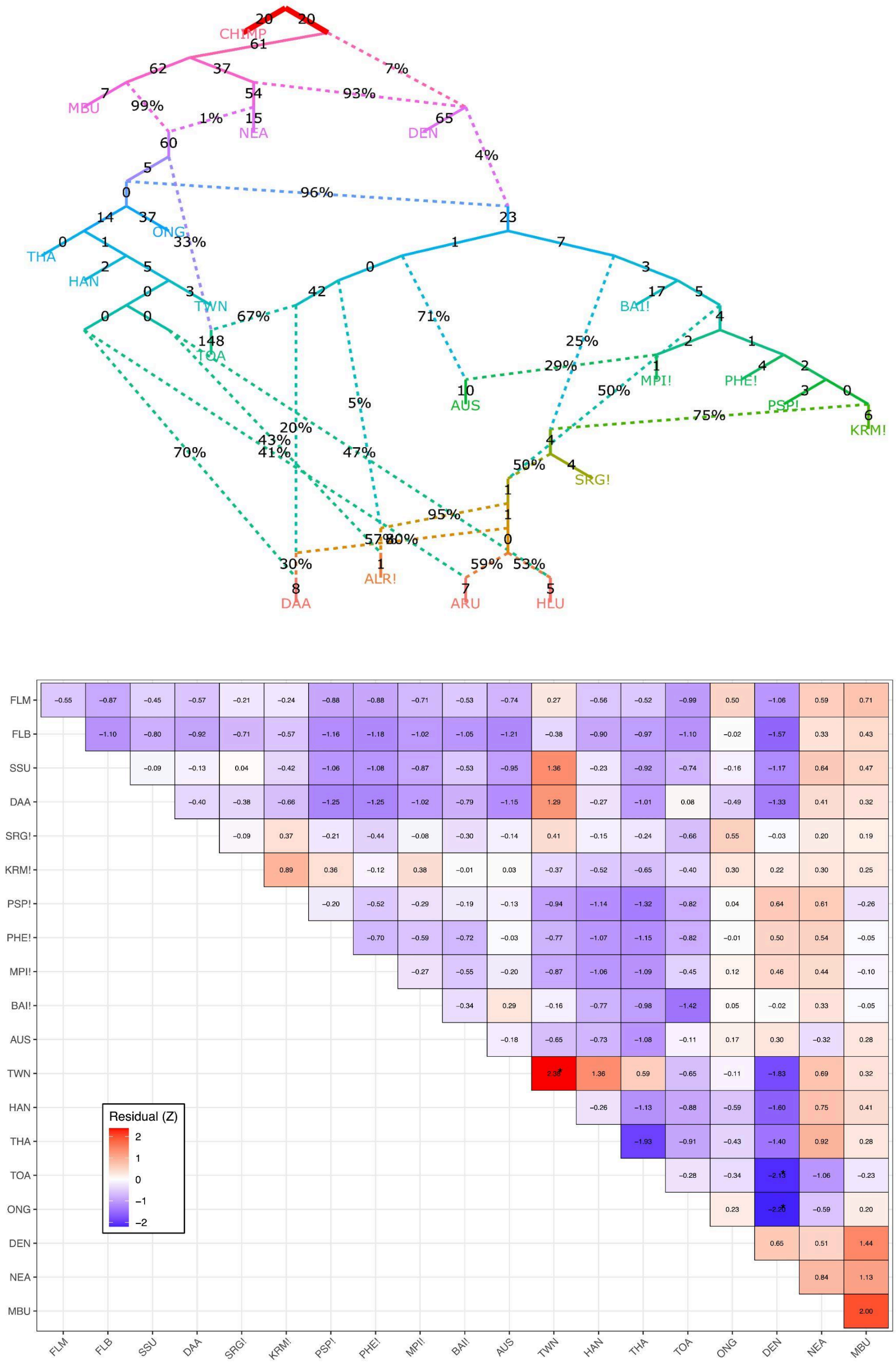


**Figure S14. Admixture graph showing Wallacean groups from all four regions, with deep AMH ancestry source removed. A.** Same admixture graph as shown in Fig. S12, but the unknown lineage that separates early from non-African groups is removed as an ancestor of modern Wallaceans. Models shown in Figs. S13 and S14 are statistically equivalent (see Table S5), suggesting that this deep lineage does not contribute to the ancestry of modern Wallaceans. **B.** All f3 residuals, |Z|, are less than 3, indicative of a well-fitted graph.

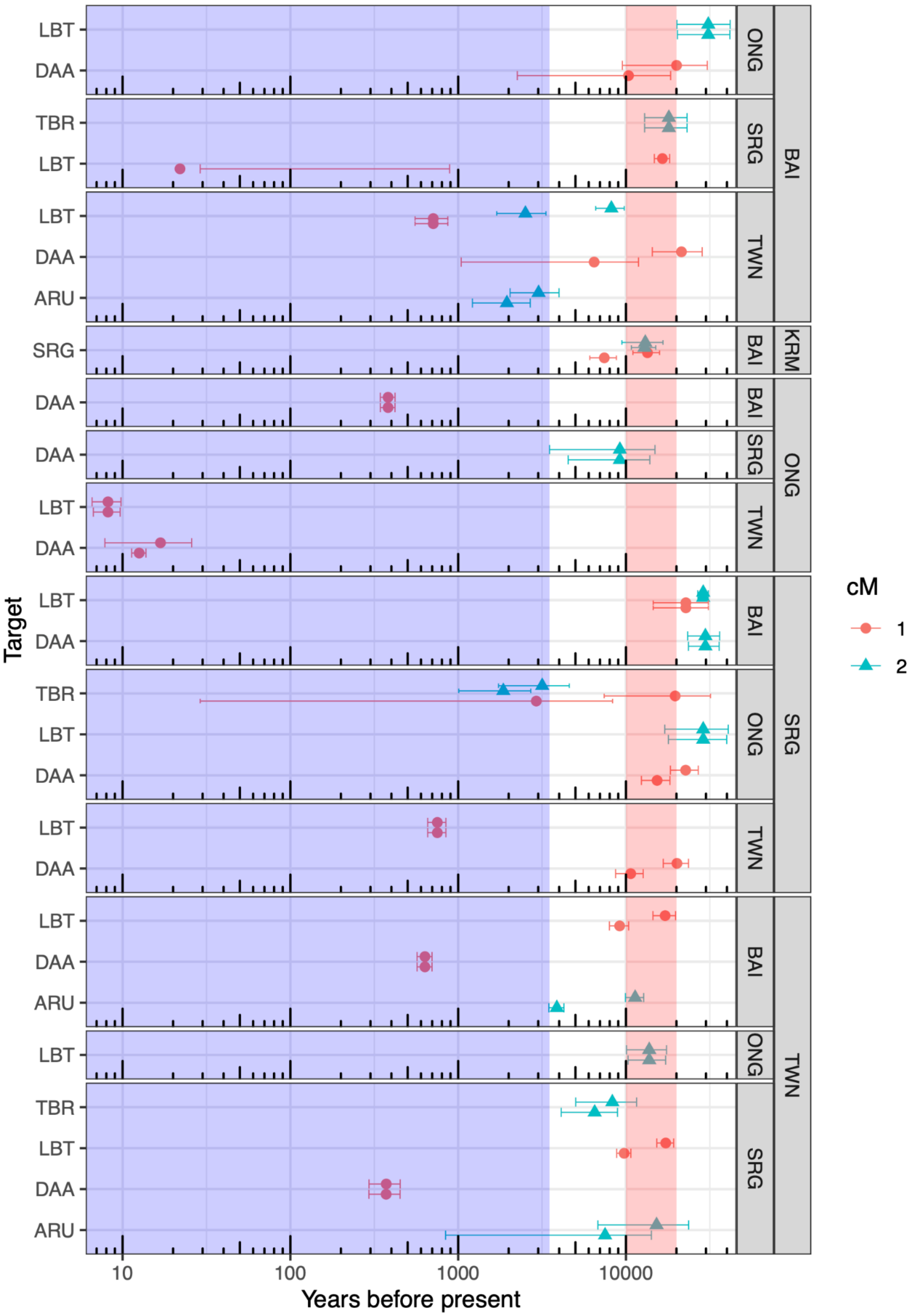

LBT
DAA
TBR
ARU
SRG
ONG
SRG
TWN
BAI
KRM
cM
1
2
Target
10
100
1000
10000
Years before present

**Figure S15. LeNata admixture estimates.** Plausible admixture times for different ancestor proxies (facet labels on right hand side of y-axis) inferred for four Wallacean (Aru, ARU; Tanimbar, TBR; Lembata, LBT; and Daa from Sulawesi, DAA)  and the West Papuan Sorong population (SRG) (Target groups, axis labels on left hand side of y-axis). LaNeta allows for two introgression pulses from one of the two ancestor groups, with the group contributing two pulses listed in the left-most facet label. Estimates are reported at different genetic intervals (1 and 2 cM; see key). Only estimates where 99% lower confidence interval exceeds time = 0 are shown. Dates postdating Austronesian arrival ~3.5 kya fall within the shaded blue area and those lying in the 10 - 20 kya interval associated with demographic change in New Guinea are indicated by the shaded red area. Notably, many dates involving an Indigenous Taiwanese group substantially predate 3.5 kya, and suggest estimation errors that may stem from a lack of samples (i.e. <10, where 10s to 100s are recommended (Liang et al. 2022)).

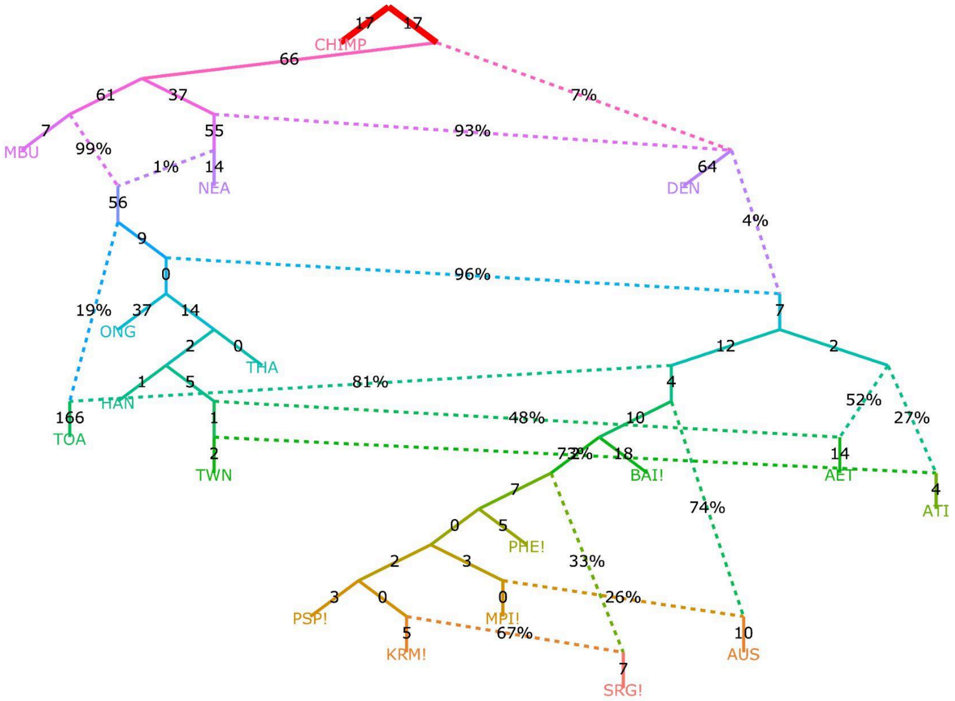

CHIMP
17
17
66
7%
61
37
55
93%
7
MBU
99%
1%
14
NEA
64
DEN
56
4%
9
0
96%
19%
37
14
ONG
7
2
0
THA
12
2
1
5
81%
4
HAN
52%
166
TOA
1
48%
10
27%
2
TWN
73%
2%
18
BAI!
14
AET
4
ATI
7
74%
0
5
PHE!
2
3
33%
3
0
0
26%
PSP!
MPI!
5
67%
10
KRM!
AUS
7
SRG!


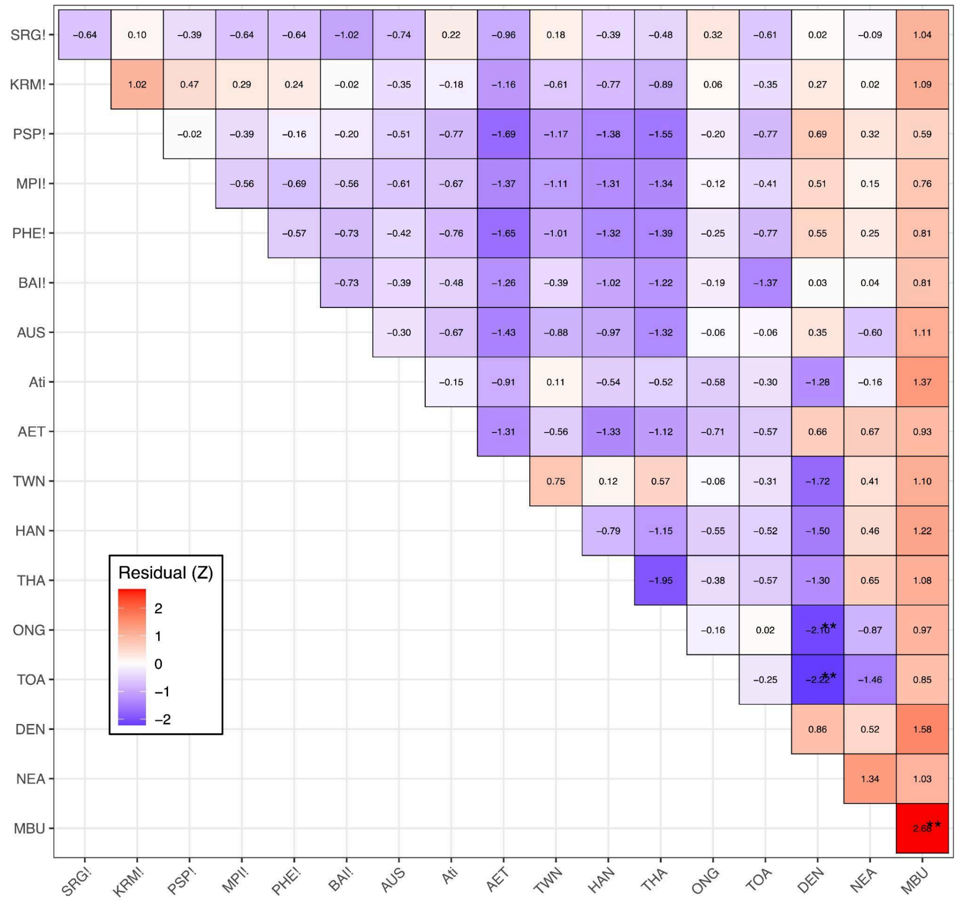

Residual (Z)
2
1
0
−1
−2
SRG! −0.64 0.10 −0.39 −0.64 −0.64 −1.02 −0.74 0.22 −0.96 0.18 −0.39 −0.48 0.32 −0.61 0.02 −0.09 1.04
KRM! 1.02 0.47 0.29 0.24 −0.02 −0.35 −0.18 −1.16 −0.61 −0.77 −0.89 0.06 −0.35 0.27 0.02 1.09
PSP! −0.02 −0.39 −0.16 −0.20 −0.51 −0.77 −1.69 −1.17 −1.38 −1.55 −0.20 −0.77 0.69 0.32 0.59
MPI! −0.56 −0.69 −0.56 −0.61 −0.67 −1.37 −1.11 −1.31 −1.34 −0.12 −0.41 0.51 0.15 0.76
PHE! −0.57 −0.73 −0.42 −0.76 −1.65 −1.01 −1.32 −1.39 −0.25 −0.77 0.55 0.25 0.81
BAI! −0.73 −0.39 −0.48 −1.26 −0.39 −1.02 −1.22 −0.19 −1.37 0.03 0.04 0.81
AUS −0.30 −0.67 −1.43 −0.88 −0.97 −1.32 −0.06 −0.06 0.35 −0.60 1.11
Ati −0.15 −0.91 0.11 −0.54 −0.52 −0.58 −0.30 −1.28 −0.16 1.37
AET −1.31 −0.56 −1.33 −1.12 −0.71 −0.57 0.66 0.67 0.93
TWN 0.75 0.12 0.57 −0.06 −0.31 −1.72 0.41 1.10
HAN −0.79 −1.15 −0.55 −0.52 −1.50 0.46 1.22
THA −1.95 −0.38 −0.57 −1.30 0.65 1.08
ONG −0.16 0.02 −2.10** −0.87 0.97
TOA −0.25 −2.22** −1.46 0.85
DEN 0.86 0.52 1.58
NEA 1.34 1.03
MBU 2.68**
SRG! KRM! PSP! MPI! PHE! BAI! AUS Ati AET TWN HAN THA ONG TOA DEN NEA MBU

**Figure S16. Admixture graph showing Philippines groups within a broad hominin context. A.** Well fitted qpGraph identified by *find_graph* algorithm with minor modifications. **B.** All f3 residuals, |Z|, are less than 3, indicative of a well-fitted graph.

Byrska-Bishop, Evani, Zhao, and Basile. 2021. “High Coverage Whole Genome Sequencing of the Expanded 1000 Genomes Project Cohort Including 602 Trios. bioRxiv.” *Published Online February*.

Chen, Shifu, Yanqing Zhou, Yaru Chen, and Jia Gu. 2018. “Fastp: An Ultra-Fast All-in-One FASTQ Preprocessor.” *Bioinformatics* 34 (17): i884–90.

Danecek, Petr, James K. Bonfield, Jennifer Liddle, John Marshall, Valeriu Ohan, Martin O. Pollard, Andrew Whitwham, et al. 2021. “Twelve Years of SAMtools and BCFtools.” *GigaScience* 10 (2). https://doi.org/10.1093/gigascience/giab008.

Harney, Éadaoin, Nick Patterson, David Reich, and John Wakeley. 2021. “Assessing the Performance of qpAdm: A Statistical Tool for Studying Population Admixture.” *Genetics* 217 (4): iyaa045.

Lan, Divon Mordechai, Gludhug Purnomo, Raymond Tobler, Yassine Souilmi, and Bastien Llamas. 2022. “Genozip Dual-Coordinate VCF Format Enables Efficient Genomic Analyses and Alleviates Liftover Limitations.” *bioRxiv*. https://doi.org/10.1101/2022.07.17.500374.

Larena, Maximilian, Federico Sanchez-Quinto, Per Sjödin, James McKenna, Carlo Ebeo, Rebecca Reyes, Ophelia Casel, et al. 2021. “Multiple Migrations to the Philippines during the Last 50,000 Years.” *Proceedings of the National Academy of Sciences of the United States of America* 118 (13). https://doi.org/10.1073/pnas.2026132118.

Liang, Mason, Mikhail Shishkin, Anastasia Mikhailova, Vladimir Shchur, and Rasmus Nielsen. 2022. “Estimating the Timing of Multiple Admixture Events Using 3-Locus Linkage Disequilibrium.” *PLoS Genetics* 18 (7): e1010281.

Li, Heng. 2013. “Aligning Sequence Reads, Clone Sequences and Assembly Contigs with BWA-MEM.” *arXiv: Genomics*. arXiv. http://arxiv.org/abs/1303.3997.

Li, Heng, Bob Handsaker, Alec Wysoker, Tim Fennell, Jue Ruan, Nils Homer, Gabor Marth, Goncalo Abecasis, Richard Durbin, and 1000 Genome Project Data Processing Subgroup. 2009. “The Sequence Alignment/Map Format and SAMtools.” *Bioinformatics* 25 (16): 2078–79.

Maier, Robert, Pavel Flegontov, Olga Flegontova, Piya Changmai, and David Reich. 2022. “On the Limits of Fitting Complex Models of Population History to Genetic Data.” *bioRxiv*. https://doi.org/10.1101/2022.05.08.491072.

Maier, Robert, Pavel Flegontov, Olga Flegontova, Ulaş Işıldak, Piya Changmai, and David Reich. 2023. “On the Limits of Fitting Complex Models of Population History to F-Statistics.” *eLife* 12 (June). https://doi.org/10.7554/eLife.85492.

Maier, Robert, and Nick Patterson. 2024. *Admixtools: Inferring Demographic History from Genetic Data*. https://github.com/uqrmaie1/admixtools.

McKenna, Aaron, Matthew Hanna, Eric Banks, Andrey Sivachenko, Kristian Cibulskis, Andrew Kernytsky, Kiran Garimella, et al. 2010. “The Genome Analysis Toolkit: A MapReduce Framework for Analyzing next-Generation DNA Sequencing Data.” *Genome Research* 20 (9): 1297–1303.

Nassar, Luis R., Galt P. Barber, Anna Benet-Pagès, Jonathan Casper, Hiram Clawson, Mark Diekhans, Clay Fischer, et al. 2023. “The UCSC Genome Browser Database: 2023 Update.” *Nucleic Acids Research* 51 (D1): D1188–95.

Oliveira, Sandra, Kathrin Nägele, Selina Carlhoff, Irina Pugach, Toetik Koesbardiati, Alexander Hübner, Matthias Meyer, et al. 2022. “Ancient Genomes from the Last Three Millennia Support Multiple Human Dispersals into Wallacea.” *Nature Ecology & Evolution* 6 (7): 1024–34.

Patterson, Nick, Alkes L. Price, and David Reich. 2006. “Population Structure and Eigenanalysis.” *PLoS Genetics* 2 (12): e190.

Price, Alkes L., Nick J. Patterson, Robert M. Plenge, Michael E. Weinblatt, Nancy A. Shadick, and David Reich. 2006. “Principal Components Analysis Corrects for Stratification in Genome-Wide Association Studies.” *Nature Genetics* 38 (8): 904–9.

Purcell, Shaun, Benjamin Neale, Kathe Todd-Brown, Lori Thomas, Manuel A. R. Ferreira, David Bender, Julian Maller, et al. 2007. “PLINK: A Tool Set for Whole-Genome Association and Population-Based Linkage Analyses.” *American Journal of Human Genetics* 81 (3): 559–75.

Rubinacci, Simone, Diogo M. Ribeiro, Robin J. Hofmeister, and Olivier Delaneau. 2021. “Efficient Phasing and Imputation of Low-Coverage Sequencing Data Using Large Reference Panels.” *Nature Genetics* 53 (1): 120–26.

Tischler, German, and Steven Leonard. 2014. “Biobambam: Tools for Read Pair Collation Based Algorithms on BAM Files.” *Source Code for Biology and Medicine* 9 (1): 13.

Yüncü, Eren, Ulaş Işıldak, Matthew P. Williams, Christian D. Huber, Leonid A. Vyazov, Piya Changmai, and Pavel Flegontov. 2023. “False Discovery Rates of qpAdm-Based Screens for Genetic Admixture.” *bioRxiv*. https://doi.org/10.1101/2023.04.25.538339.
